# Wide distribution of phage that infect freshwater SAR11 bacteria

**DOI:** 10.1101/672428

**Authors:** Lin-Xing Chen, Yan-Lin Zhao, Katherine D. McMahon, Jiro F. Mori, Gerdhard L. Jessen, Tara Colenbrander Nelson, Lesley A. Warren, Jillian F. Banfield

**Author notes:** Corresponding author: Jillian F. Banfield, Address: McCone Hall, Berkeley, CA 94720.

## Abstract

*Fonsibacter* (LD12 subclade) are among the most abundant bacterioplankton in freshwater ecosystems. These bacteria belong to the order Pelagibacterales (SAR11) and are related to *Pelagibacter* (marine SAR11) that dominate many marine habitats. Although a handful of *Pelagibacter* phage (Pelagiphage) have been described, no phage that infect *Fonsibacter* have been reported. In this study, a complete *Fonsibacter* genome containing a prophage was reconstructed from metagenomic data. A circularized and complete genome related to the prophage, referred to as uv-Fonsiphage-EPL, shows high similarity to marine Pelagiphage HTVC025P. Additionally, we reconstructed three complete and one draft genome of phage related to marine Pelagiphage HTVC010P, and predicted a lytic strategy. The similarity in codon usage and co-occurrence patterns of HTVC010P-related phage and *Fonsibacter* suggested that these phage infect *Fonsibacter*. Similar phage were detected in Lake Mendota, Wisconsin, where *Fonsibacter* is also present. A search of related phage revealed the worldwide distribution of some genotypes in freshwater ecosystems, suggesting their substantial role in shaping indigenous microbial assemblages and influence on biogeochemical cycling. However, the uv-Fonsiphage-EPL and one lineage of HTVC010P-related phage have a more limited distribution in freshwater ecosystems. Based on this, and their close phylogenetic relatedness with Pelagiphage, we predict that they transitioned from saline into freshwater ecosystems comparatively recently. Overall, the findings provide insights into the genomic features of phage that infect *Fonsibacter*, and expand understanding of the ecology and evolution of these important bacteria.

## Introduction

Heterotrophic SAR11 bacteria (Alphaproteobacteria; Pelagibacterales) are often very abundant in marine and freshwater ecosystems (Salcher et al. 2011; Grote et al. 2012; Zaremba-Niedzwiedzka et al. 2013; Cabello-Yeves et al. 2018). *Fonsibacter*, the freshwater subclade of SAR11, also known as LD12 (or III-b), are especially abundant in the euphotic layers of lakes during summer (Salcher et al. 2011), and play important roles in the assimilation of low molecular weight carboxylic acids (Salcher et al. 2011; Eiler et al. 2016). The first *Fonsibacter* genomes were reconstructed via single-cell genomics and subsequent analyses indicated their low recombination rates in nature (Zaremba-Niedzwiedzka et al. 2013). Comparative genomic analyses showed many proteins shared between *Fonsibacter* and *Pelagibacter* (marine SAR11), but metabolic divergence was also detected (Grote et al. 2012). *Fonsibacter* typically use the Embden–Meyerhof–Parnass (EMP) rather than the Entner–Doudoroff glycolysis pathway, and produce rather than uptake osmolytes (Dupont et al. 2014; Eiler et al. 2016). These studies proposed that *Fonsibacter* evolved from a streamlined ancestor of marine *Pelagibacter* (Zaremba-Niedzwiedzka et al. 2013; Eiler et al. 2016). It was initially proposed that the transition between marine and freshwater ecosystems happened only once (Logares et al. 2010), but this conclusion was challenged recently. For example, a metagenomics-assembled genome from the freshwater Lake Baikal was phylogenetically assigned to *Pelagibacter* (Cabello-Yeves et al. 2018), and phylogenetic analyses of 16S rRNA genes suggested the existence of several marine SAR11 subtypes in freshwater lakes (Paver et al. 2018). The first cultivated representative of *Fonsibacter* isolated from southern Louisiana coast (Henson et al. 2018), reported very recently, has isocitrate lyase for a complete glyoxylate bypass of the TCA cycle along with malate synthase, distinguishing it from other *Fonsibacter.* The authors suggest temperature-based ecotype diversification within this genus.

SAR11 rarely use CRISPR-Cas or restriction-modification systems for phage defense (Zaremba-Niedzwiedzka et al. 2013; Zhao et al. 2013; Giovannoni 2017). However, they harbor the hypervariable region 2 located between their 16S/23S rRNA and 5S rRNA genes, which contains genes encoding various transferases, isomerases, O-antigen and pilins. SAR11 may use these proteins to defend against phage by cell surface modification (Grote et al. 2012; Zaremba-Niedzwiedzka et al. 2013; Henson et al. 2018). To date, 15 Pelagibacter phage (Pelagiphages) have been isolated from marine environments (Zhao et al. 2013; Zhao et al. 2018) and Pelagiphage HTVC010P is suggested to be among the most abundant phage in the ocean (Zhao et al. 2013). In contrast, no phage that infect *Fonsibacter* have been reported. In general, phage that infect major heterotrophic groups in freshwater ecosystems are largely unknown, with only a few cases reported recently, including phage of LD28 clade (‘*Ca*. Methylopumilus planktonicus’) (Moon et al. 2017) and the Actinobacteria acl clade (Ghai et al. 2017). Phage that infect freshwater heterotrophic bacteria groups could shape the freshwater microbial assemblages and redistribute bacterially-derived compounds via the lysis of host cells. Thus, phage of heterotrophic freshwater bacteria may significantly influence biogeochemical cycles, especially of carbon.

Here, we performed genome-resolved metagenomic analyses on microbial communities from freshwater ecosystems to reconstruct genomes of *Fonsibacter* and their phage. Comparative analyses of *Fonsibacter* and *Pelagibacter* infecting phage show genetic conservation and divergence. The distribution of some related phage in freshwater ecosystems suggests the broad ecological significance of *Fonsibacter* phage. Overall, the findings shed light on the ecology of *Fonsibacter*, and reveal aspects of phage and host evolutionary history.

## Results

### Metagenome-assembled genome of Fonsibacter

Freshwater samples were collected from an End Pit Lake (EPL) in Alberta, Canada (methods). Analysis of the EPL metagenomic datasets (Supplementary Table 1) revealed one genome bin with 27 scaffolds, two of which had features indicative of a prophage. Subsequently, this bin was manually curated into a complete genome. The genome accuracy was verified based on paired read mapping throughout. It contains no repeats long enough to have confounded the assembly and displays GC skew and cumulative GC skew with the form expected for complete bacterial genomes that undergo bidirectional replication (Supplementary Fig.1). The genome is 1,136,868 bp in length, the smallest SAR11 genome yet reported, and has a GC content of 29.6% (Table 1). Phylogenetic analyses based on a set of 16 ribosomal proteins indicated the genomically defined bacterium belongs to the candidate genus *Fonsibacter* (Fig.1a). The 16S rRNA gene sequence of this *Fonsibacter* genome is identical to that of AAA028-C07 recovered from Lake Mendota, Wisconsin (Zaremba-Niedzwiedzka et al. 2013) and shares 99.8% identity with that of the *Fonsibacter* isolate *Candidatus* Fonsibacter ubiquis LSUCC0530 (Henson et al. 2018). The new *Fonsibacter* genome shares 96% and 86% genome-wide average nucleotide identity (ANI) with the 0.85 Mbp AAA028-C07 draft and 1.16 Mbp complete LSUCC0530 genomes, respectively. We refer to the newly described complete genome as “EPL_02132018_0.5m_Candidatus_Fonsibacter_30_26” (hereafter “*Fonsibacter*_30_26”).

**Fig.1.**
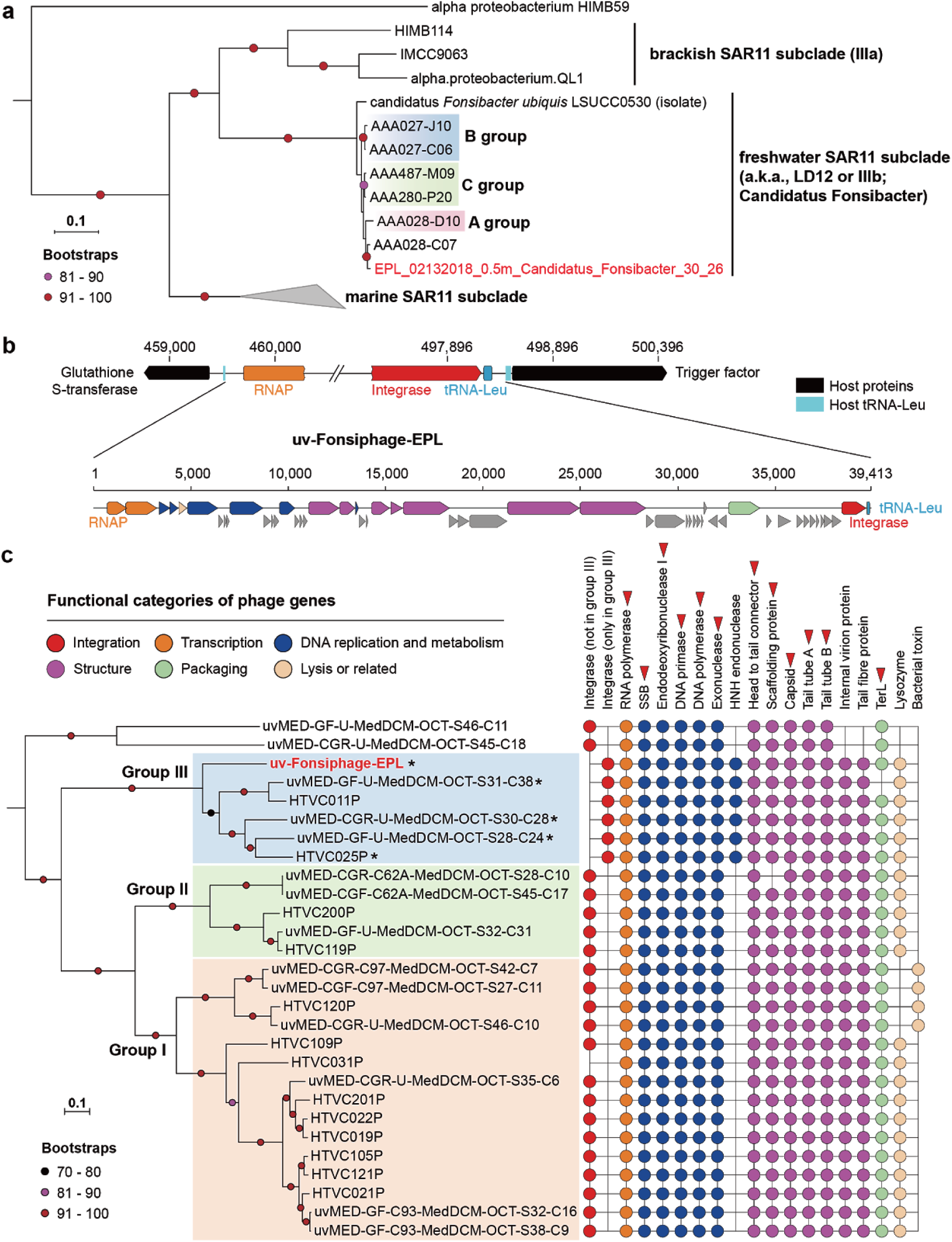
The complete *Fonsibacter* genome and its prophage. (a) Phylogenetic analyses of the complete *Fonsibacter* genome based on 16 ribosomal proteins (Methods). The three *Fonsibacter* groups defined previously are shown. The tree was rooted using the HIMB59 sequence. (b) The prophage of the complete *Fonsibacter* genome. The insertion site of the phage genome into the host tRNA-Leu is shown. Refer to (c) for the colors of different functional categories, and hypothetical proteins are indicated in grey. (c) Phylogenetic analyses of uv-Fonsiphage-EPL and related phage based on 12 core proteins (red triangles; Methods), the three *HTVC019Pvirus* groups defined recently are shown. The presence of protein families with predicted function in the phage is shown on the right (Supplementary Table 3). The phage with a fragmented RNA polymerase is indicated by an asterisk. SSB, Single-stranded DNA-binding protein.

**Table 1.** General features of the Fonsibacter and infecting phage genomes reconstructed in this study. The draft genome of an HTVC010P-related phage from BMMRE (methods) is also included, which could not be closed due to low sequencing coverage. The predicted life strategy and the closest reference for each phage is shown. Please note that the *Fonsibacter* genome lacks one of the 51 SCGs used for completeness evaluation, see main text and Supplementary Information for details. * Complete genome. ^#^ Draft genome.

| Genome<br>(short name in the main text if provided) | Life strategy | Related to | Length (bp) | GC content (%) | Completeness (%) |  | Number of |  |  |
| --- | --- | --- | --- | --- | --- | --- | --- | --- | --- |
|  |  |  |  |  | 51 SCGs | CheckM | rRNAs | tRNAs | Proteins |
| EPL_02132018_0.5m_Candidatus_Fonsibacter_30_26 *<br>(Fonsibacter_30_26) | — | AAA028-C07 | 1,136,868 | 29.6 | 50 | 100 | 3 (5S,16S,23S) | 31 | 1,229 |
| uv-Fonsiphage-EPL * | Lysogenic | Pelagiphage HTVC025P | 39,413 | 32.1 | — | — | — | 1 | 52 |
| EPL_06132017_6.25m_HTVC010P-related_33_76 *<br>(HTVC010P-related_33_76) | Lytic | Pelagiphage HTVC010P | 35,816 | 32.5 | — | — | — | — | 61 |
| EPL_08022017_1.5m_HTVC010P-related_32_16 *<br>(HTVC010P-related_32_16) | Lytic | Pelagiphage HTVC010P | 36,457 | 31.9 | — | — | — | — | 60 |
| I-EPL_09192017_0.5m_HTVC010P-related_33_10 *<br>(HTVC010P-related_33_10) | Lytic | Pelagiphage HTVC010P | 36,507 | 32.5 | — | — | — | — | 62 |
| BMMRE_07242016_10_scaffold_124 # | Lytic | Pelagiphage HTVC010P | 27,140 | 31.6 | — | — | — | — | 33 |

The *Fonsibacter*_30_26 genome encodes 1229 protein-coding genes, 3 rRNA genes (one copy each of the 5S, 16S and 23S rRNA genes) and 31 tRNAs (Table 1). *Fonsibacter*_30_26 does not encode the gene for the 50S ribosomal protein L30, a feature we predict is shared by all reported SAR11 genomes (Supplementary Fig.2). *Fonsibacter*_30_26 has the full EMP glycolysis pathway, a complete gluconeogenesis pathway and a full TCA cycle, also a complete oxidative phosphorylation pathway, as reported for other *Fonsibacter* genomes (Grote et al. 2012; Zaremba-Niedzwiedzka et al. 2013; Henson et al. 2018). No carbon fixation gene or pathway was identified in the genome, indicating a heterotrophic lifestyle of this *Fonsibacter* species. Interestingly, within a region previously described to be hypervariable in SAR11 (53,690 bp in length, 54 protein-coding genes; Supplementary Figs.3 and 4), we detected four genes encoding transketolase, one of the three enzymes in the non-oxidative pentose phosphate pathway. However, these transketolases only contained one or two of the three domains found in a full-length transketolase sequence, therefore their function in the pentose phosphate pathway remains uncertain. We identified 18 genes in the hypervariable region that encode glycosyltransferase, methyltransferase and epimerase, which are common in SAR11 and may be involved in phage defense (Grote et al. 2012; Zaremba-Niedzwiedzka et al. 2013; Henson et al. 2018).

### The first genome of phage infecting *Fonsibacter*

We mapped metagenomic reads to the putative *Fonsibacter*_30_26 prophage region and recovered reads that could be reconstructed into complete phage genome (Supplementary Information). The prophage genome is inserted between *attL* (left end of prophage) and *attR* (right end of prophage), which share an 11 bp identical ‘core sequence’. Specifically, ∼5% of reads circularized the phage genome, indicating the presence of some free phage particles. In addition, some bacterial cells lack the prophage, so the prophage start and end could be clearly defined. We refer to the reconstructed sequence as uv-Fonsiphage-EPL. To our knowledge, this is the first genome of phage infecting *Fonsibacter*.

The genome of uv-Fonsiphage-EPL has a length of 39,413 bp and GC content of 32.1%, and encodes 52 proteins (Table 1), including integrase, DNA metabolism and replication genes, phage structural genes, lysis gene and large terminase (TerL) (Fig.1b). A search of the TerL sequence against the NCBI database revealed that uv-Fonsiphage-EPL is most closely related to phage from marine habitats. This was confirmed using phylogenetic analyses based on 12 core phage proteins (Fig.1c, Supplementary Tables 2 and 3). In detail, uv-Fonsiphage-EPL grouped with two Pelagiphage isolates from Baltic sea (HTVC025P) (Zhao et al. 2018) and Oregon coast seawater (HTVC011P) (Zhao et al. 2013), and three metagenomically-retrieved phage from the Mediterranean (Mizuno et al. 2013), within the *HTVC019Pvirus* (*Caudovirales*, *Podoviridae*, *Autographivirinae*) group III defined recently (Zhao et al. 2018). The uv-Fonsiphage-EPL shares 75.8-78.5% TerL similarity with group III members and most similar to HTVC025P. Genome-wide alignment revealed conserved genome synteny and high similarity between The uv-Fonsiphage-EPL and two Pelagiphage isolates (Supplementary Fig.5a).

The phage in the *HTVC019Pvirus* group III share several features (Fig.1c). For example, they have an integrase with limited degree of identity with those in group I and II members, and an HNH endonuclease that is absent in the other two groups. Although the integrases shared low similarities within group III (31.6-36.9%), uv-Fonsiphage-EPL, HTVC011P and HTVC025P all can integrate into the host tRNA-Leu (TAG) site, and the ‘core sequence’ (Supplementary Information) in uv-Fonsiphage-EPL is only one base pair different from that of HTVC025P(Zhao et al. 2018). The phage encodes its own tRNA-Leu, replacing the lost function of the host tRNA-Leu gene after phage integration (Fig.1b). A bacterial trigger factor protein flanks the prophage in all three host genomes (Fig.1b and ref (Zhao et al. 2018)). Also, divergence was detected among *HTVC019Pvirus* group III members, for example, uv-Fonsiphage-EPL lacked several hypothetical proteins found in most of the other phage (Supplementary Table 3).

Interestingly, all of the RNA polymerase genes in all group III genomes, except HTVC011P, were fragmented into two parts (Fig.1c). We did not identify any RNA polymerase reads mapped to the uv-Fonsiphage-EPL genome that were not split. This, in combination with detection of the split gene in the other genomes, suggests that gene interruption did not occur recently.

### Metagenome-assembled genomes of potential *Fonsibacter-*infecting phage

Given the high similarity among uv-Fonsiphage-EPL and the *HTVC019Pvirus* Pelagiphage (see above), we expected to detect counterparts of other type of marine Pelagiphage in freshwater ecosystems (Zhao et al. 2013). A subset of scaffolds from samples of EPL, I-EPL (the input source of EPL) and BMMRE, a Base Metal Mine Receiving Environment in Manitoba of Canada (methods), encode TerL that share 62-85% amino acid with that of Pelagiphage HTVC010P (*Podoviridae*). Manual curation generated three distinct complete genomes and one draft genome. These are referred to as HTVC010P-related phage (Table 1, Fig.2a), and showed genome wide similarity with HTVC010P (Supplementary Fig.5b).

**Fig.2.**
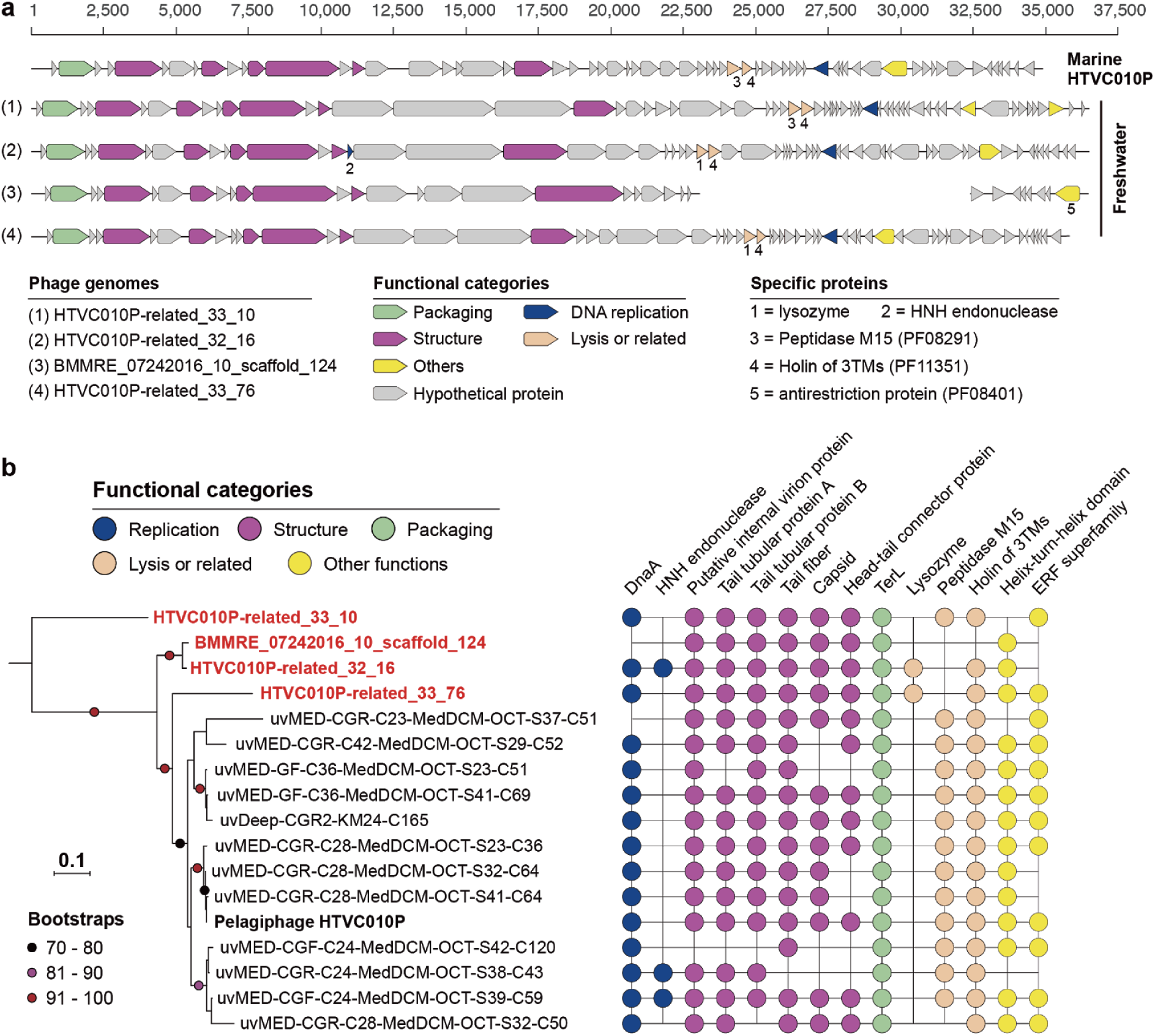
Pelagiphage HTVC010P-related phage genomes reconstructed in this study. (a) Gene content of the four phage genomes (three complete and one draft), compared to that of the marine Pelagiphage HTVC010P. Genes with predicted annotations are marked with different colors according to their function. The scaffold representing the BMMRE draft genome was split into two for better visual comparison of the genes. (b) Phylogenetic analyses of phage from this study (in red and bold) and those from marine environments based on the TerL. For each genome, the presence of protein families with predicted function in the phage is shown on the right (Supplementary Table 4).

We identified phage-specific proteins in all the HTVC010P-related genomes, including dnaA, TerL, internal protein A, tail tubular protein A and B, tail fiber, capsid, head-tail connector protein (Fig.2, Supplementary Table 4). No integrase was detected, suggesting that they are lytic phages. An HNH endonuclease was identified in HTVC010P-related_32_16 and two marine phage genomes but not in the BMMRE draft genome, though they are phylogenetically closely related (Fig.2b). Interestingly, within all the HTVC010P-related genomes, only two of those reconstructed in this study harbored a lysozyme protein. Instead, those without lysozyme may use a peptidase M15 (PF08291) for the cell lysis function (Supplementary Information). No lysozyme or peptidase M15 was detected in the BMMRE draft genome, likely due to incompleteness. It is the only phage genome analyzed here that encodes a putative antirestriction protein (PF08401), possibly for protecting its DNA against host endonuclease activity. The HTVC010P-related phage genomes also shared genes encoding many hypothetical proteins (Supplementary Table 4), suggesting their potentially important function.

### Evidence that HTVC010P-related phage infect *Fonsibacter*

We speculated that the lytic HTVC010P-related phage could infect *Fonsibacter*, given their close relationship with Pelagiphage HTVC010P. Matches of spacers from CRISPR-Cas systems to the phage genome (Andersson & Banfield 2008) and similar tRNA sequence (Bailly-Bechet et al. 2007) can be used to predict phage-host associations. Unfortunately, no CRISPR-Cas system was detected in *Fonsibacter* related scaffolds in the EPL/I-EPL/BMMRE samples. Further, we did not detect any phylogenetically informative host-associated genes in the phage genomes that could indicate host range.

Phage use host translational mechanisms during their lytic cycle (Salmond & Fineran 2015), so they may adapt to host-preferred codons (Lucks et al. 2008; Lajoie et al. 2013; Ivanova et al. 2014). Thus, codon usage bias is another approach to infer host-phage associations. We clustered all bacterial, archaeal and the four HTVC010P-related phage genomes from the same samples based on their codon usage frequency (Supplementary Table 5). The results showed that the four phage clustered with all 13 *Fonsibacter* genomes (Fig.3a), along with two Gammaproteobacteria and one Bacteroidetes genomes.

**Fig.3.**
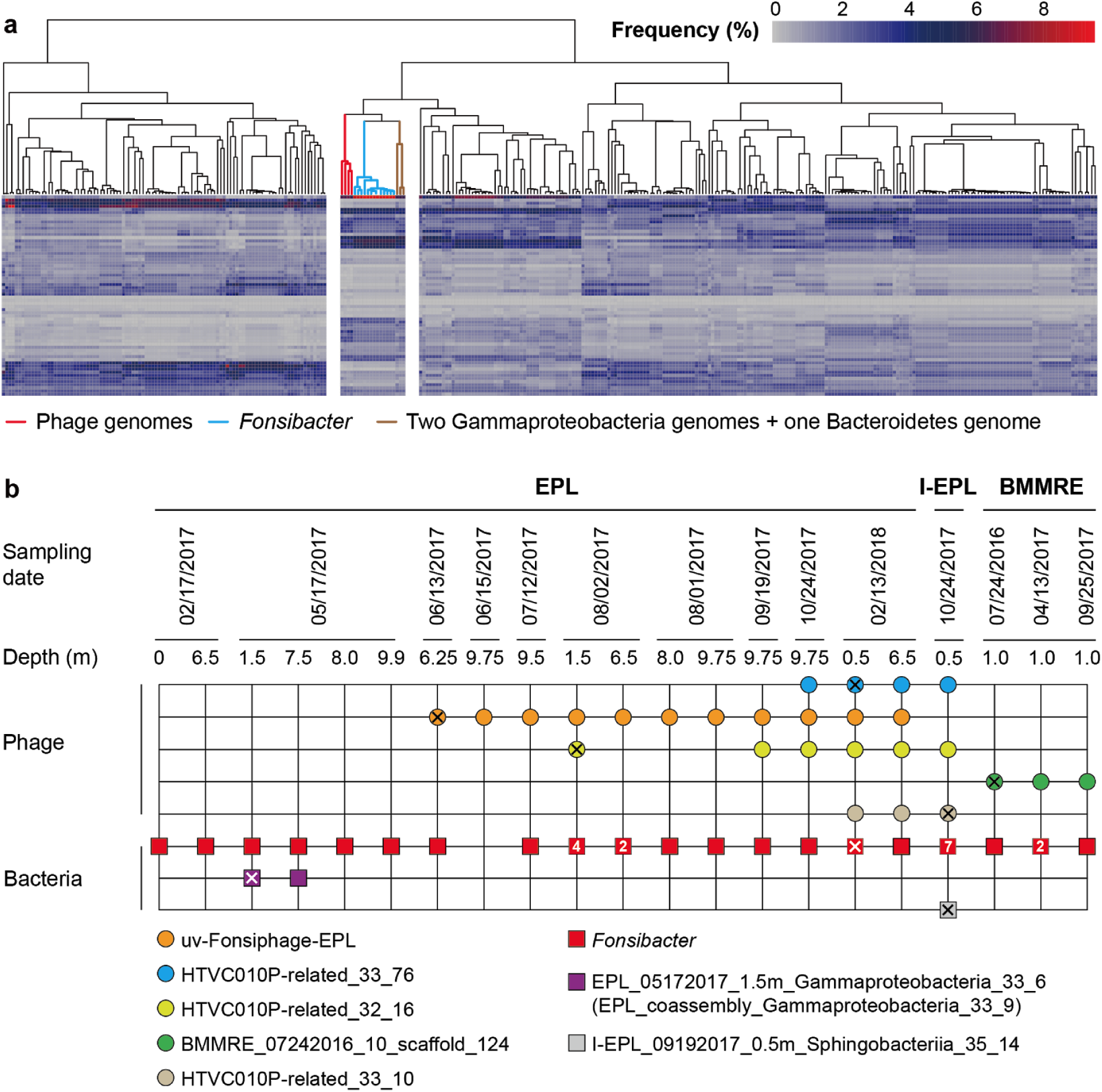
Evidence supporting the infection of *Fonsibacter* by HTVC010-related phage. (a) Clustering of genomes based on codon usage frequency. Each column represents a genome, and each line represents a codon type. The subclusters, including phage, *Fonsibacter* and three other genomes, are separate. (b) The occurrence of phage (circles), and *Fonsibacter* and the three potential host bacteria (squares). All genotypes were detected once unless a number is given inside the circle or square. The sample from where the phage or bacterial genomes were reconstructed is indicated by “X”.

We evaluated the co-occurrence of *Fonsibacter,* the two Gammaproteobacteria and one Bacteroidetes species and the phage in the EPL/I-EPL/BMMRE samples, and found that all but one of the samples that contained HTVC010P-related phage had at least one *Fonsibacter* genotype (Fig.3b, Supplementary Fig.6). However, of the samples that contained HTVC010P-related phage, the Bacteroidetes was detected in only the I-EPL sample. The two Gammaproteobacteria were detected in only two samples, neither of which contained the phage (Fig.3b). In combination, the co-occurrence patterns strongly support the inference that the HTVC010P-related phage infect *Fonsibacter* bacteria. Some samples contained only one *Fonsibacter type* and multiple phage genotypes (e.g., EPL_02/13/2018_6.5m; Fig.3b), and some samples contained only one phage but multiple *Fonsibacter* types (e.g., BMMRE_04/13/2017_1.0m; Fig.3b). These findings indicate the “multiple vs. multiple” host-phage relationship, in line with previous studies on *Pelagibacter* and its phage (Mizuno et al. 2013; Zhao et al. 2013; Zhao et al. 2018).

### *Fonsibacter* and their phage in Lake Mendota

To further investigate the potential distribution of *Fonsibacter* and their phage, we analyzed a time series metagenomic dataset from Lake Mendota, where the first *Fonsibacter* genomes were reported (Zaremba-Niedzwiedzka et al. 2013). For this site, *Fonsibacter* strain dynamics were investigated over a five-year period (Garcia et al. 2018). A homolog search for TerL detected 19 HTVC010P-related TerL in 14 of the 90 Lake Mendota samples (Fig.4a, Supplementary Table 6). Eighteen of the TerL shared ≥ 97% amino acid identity with sequences from HTVC010P-related_32_16 and HTVC010P-related_33_10. One TerL sequence had a 90% similarity to that of uv-Fonsiphage-EPL. *Fonsibacter* was detected in all 90 samples, and showed an average rpS3 similarity of 99.6% to those from EPL/I-EPL/BMMRE samples (Fig.4b). The *Fonsibacter* members accounted for 2.1-24.8% (10.8% on average; Supplementary Table 7) of the bacterial communities (Fig.4c), indicating they were an important fraction of the indigenous microbiome. However, only 16 samples had a total phage relative abundance of ≥ 1% (two from 2010, the others from 2014), and up to 14.26% in the sample from October 22, 2012 (Fig.4c, Supplementary Table 7). There was no discernable pattern to explain the high Fonsiphage abundance in some samples.

**Fig.4.**
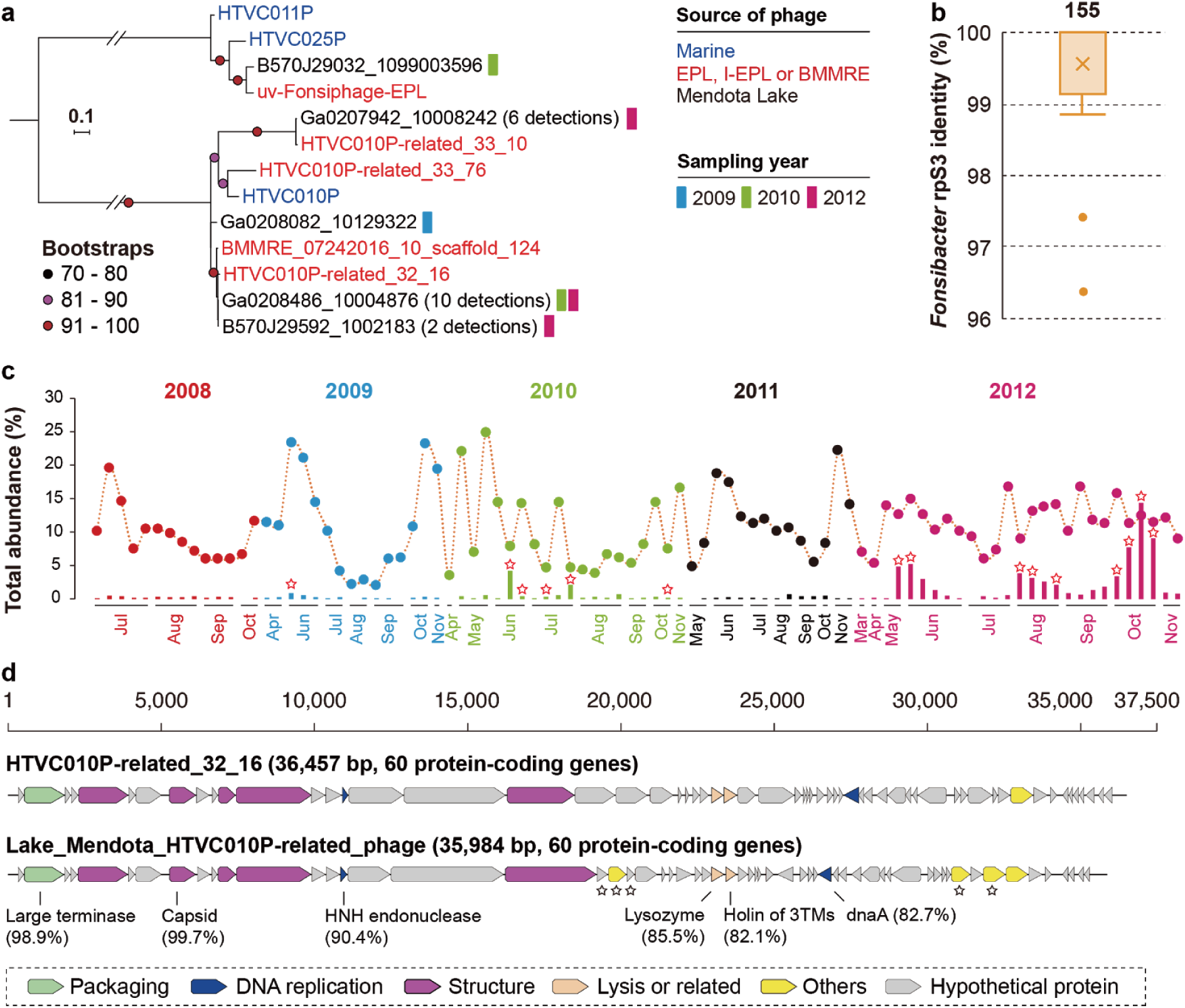
The occurrence of *Fonsibacter* and infecting phage in Lake Mendota. (a) Phylogenetic analyses of detected phage related to those from EPL/I-EPL/BMMRE in Mendota Lake based on the TerL protein. The number of genotypes is indicated in the brackets. Phage from marine habitats and reconstructed in this study were included for reference. (b) The similarity of *Fonsibacter* rpS3 proteins between EPL/I-EPL/BMMRE and Lake Mendota. The total number of *Fonsibacter* rpS3 is shown above the box plot. (c) The relative abundance of *Fonsibacter* (colored circles) and phage (colored bars) across the five-year sampling period in Lake Mendota. A red star indicates the detection of assembled TerL in the corresponding metagenomic dataset. (d) Comparative analyses of the phage genomes reconstructed from Lake Mendota (close to HTVC010P-related_31_16) and EPL (HTVC010P-related_31_16). The sequence similarities of some proteins between these two genomes are shown. A black star indicates genes not in EPL phage genome.

In spite of their high TerL similarity, Lake Mendota phage showed different genomic features from EPL/I-EPL/BMMRE HTVC010P-related phage (Supplementary Fig.7a). We reconstructed a complete 35,984 bp phage genome from the Lake Mendota datasets that encodes 60 protein-coding genes (Fig.4d, Supplementary Fig.7b). Its TerL and major capsid proteins share 98.9% and 99.7% similarity to that of HTVC010P-related_32_16. This genome of the Lake Mendota phage has several genes not found in the EPL genome (Fig.4d). It contained an HNH endonuclease that is present in HTVC010P-related_32_16 but absent in most HTVC010P-related phage (Fig.2b). Phylogenetic analyses showed these HTVC010P-related HNH endonucleases were closely related to those from uv-Fonsiphage-EPL and related phage (Supplementary Fig.8). However, we cannot distinguish whether the endonuclease was ancestral and lost in some members from acquisition via horizontal transfer.

### Wide distribution of *Fonsibacter-*infecting phage

By searching public metagenomic datasets (methods), we retrieved 403 TerL sequences from 193 freshwater-related samples and 2393 TerL sequences from 568 marine/saline samples (all shared ≥ 80% similarity to those of phage reported in this study; Supplementary Fig.9a). Overall, the freshwater-related TerL sequences shared on average 96% amino acid identity with those reported here, whereas marine/saline predicted proteins only shared on average 83% identity. However, some anomalously similar TerL sequences were detected in both habitat types (Supplementary Table 8). Eleven of the freshwater-related outliers were from Africa inland freshwater lakes, including Kabuno Bay, Lake Kivu and Lake Malawi, which are all geographically connected by the Rusizi River. Some of these phage cluster together in an Africa-specific group (lineage 2a; Fig.5), and apparently associate with an Africa-specific lineage of *Fonsibacter* (Supplementary Fig.11). The other two freshwater-related outliers were from the Alfacada pond (Ebro Delta, Spain) and associated with the skin of a European eel (Carda-Diéguez et al. 2014; Carda-Diéguez et al. 2015; Carda-Diéguez et al. 2017). For the marine/saline outliers, the majority were from San Francisco Bay, Columbia River estuary and Delaware River and Bay, which are characterized by salinity gradients, and thus could provide niches for *Fonsibacter* (see below).

**Fig.5.**
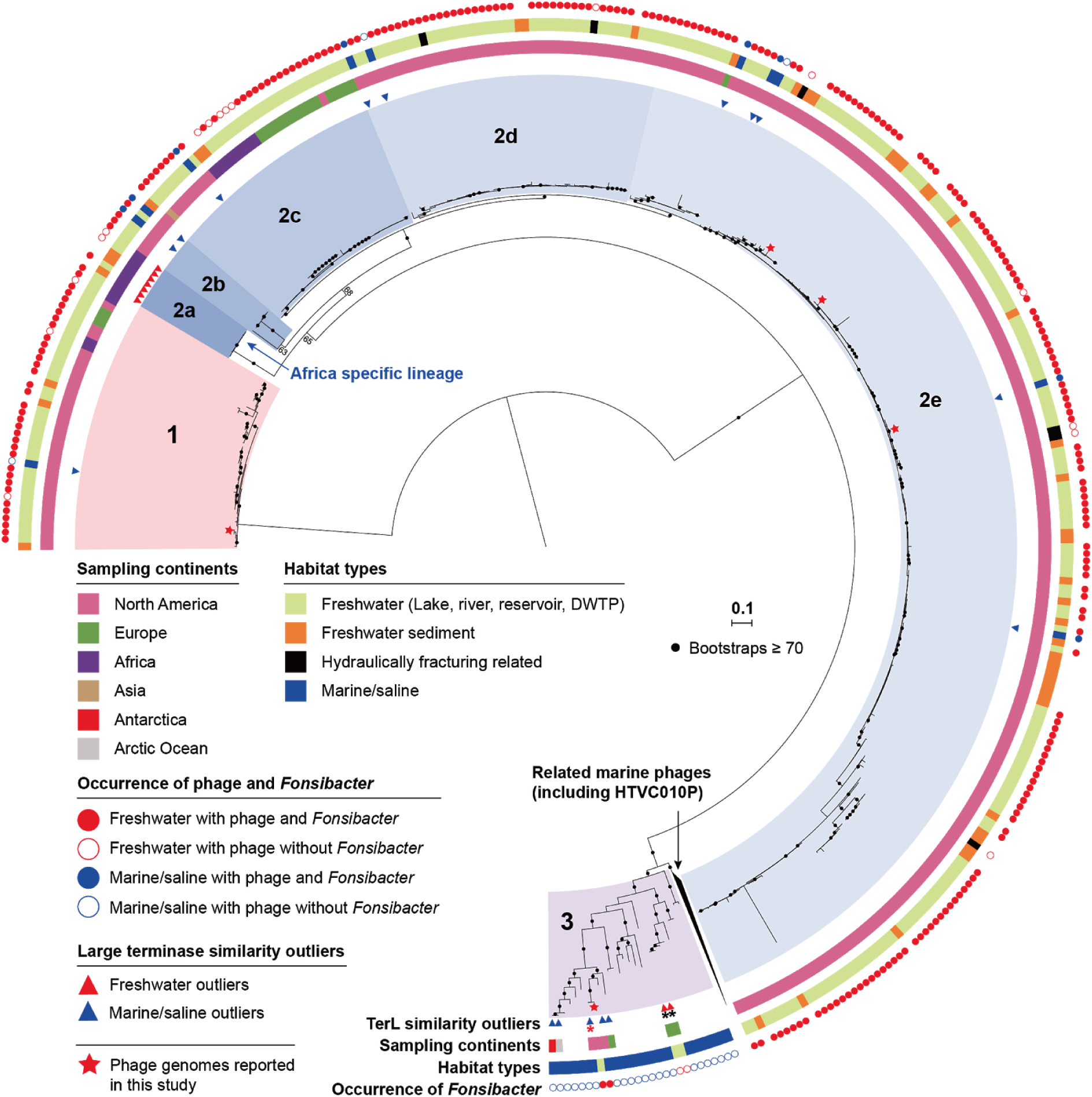
Phylogenetic analyses of HTVC010P-related phage in global freshwater ecosystems based on TerL. For phage from marine/saline habitats, only those with anomalously high similarity to the EPL/I-EPL/BMMRE phage were included, along with similar phage from marine lineage 3 as references. The similarity outliers from freshwater and related habitats, and marine/saline habitats are indicated by red and blue triangles, respectively. The TerL were assigned to habitat types and sampling continents based on sampling information. Stars represented phage whose genomes are reported in the current study. The phage were grouped into 3 lineages based on the phylogeny and TerL identity (≥ 80%), sub-lineages were determined in lineage 2 based on phylogeny. The presence/absence of *Fonsibacter* in the sample with TerL detected is indicated by solid/open circles, not shown for sediment samples because none with *Fonsibacter* detected. The TerL detected in European eel related samples are indicated by black asterisks, the one from Lake Walker sediment is indicated by a red asterisk (in lineage 3). Bootstrap values are indicated by black dots if values are ≥ 70). DWTP, drinking water treatment plant.

We investigated the distribution of *Fonsibacter* phage in freshwater-related ecosystems using detection of the TerL protein sequence. The phage were detected in 118 lake/pond/reservoir, 35 river, 34 sediment, 4 hydraulically fracturing related, and two drinking water treatment plant samples (Supplementary Fig.9b and Table 8). *Fonsibacter* was found in most samples, but not in the freshwater sediment and three hydraulically fracturing related samples (Supplementary Fig.9b). Phage related to HTVC010P-related_32_16 (lineages 2a-e; Fig.5) and HTVC010P-related_33_10 (lineage 1; Fig.5) were widely distributed. However, HTVC010P-related_33_76 (lineage 3; Fig.5) and uv-Fonsiphage-EPL phage were detected in only two and six freshwater habitats (Supplementary Fig.10), respectively. With the exception of our study and the two European eel associated samples, the HTVC010P-related_33_76 phage lineage was never detected in other freshwater-related habitats (Fig.5; Supplementary Table 8).

## Discussion

### *Fonsibacter* phage are widely distributed, but show regional diversification

We reconstructed genomes of one temperate and five lytic Fonsibacter phage (Table 1, Fig.4d) that are very similar to those of some Pelagiphage (Figs.1, 2 and 4d). Among them, uv-Fonsiphage-EPL is the only reported prophage of SAR11 so far, the detection of lysis related gene indicates that it could affect the infected *Fonsibacter* population. Given that uv-Fonsiphage-EPL lacks certain hypothetical proteins found in most *HTVC019Pvirus* group III phage, we conclude that these proteins are not necessary for infection or replication in *Fonsibacter*.

Most of the lytic HTVC010P-related phage were widely distributed, but show evidence of regional diversification (Fig.5). For example, lineages 1 and 2c were detected in at least 3 continents. Phage from the same continent tend to be more closely phylogenetically related. This suggests the existence of barriers that inhibit dispersal of most lineages, possibly related to variations in indigenous phage resistance. Alternatively, the current distribution patterns of lineages 1 and 2c may reflect several recent, independent transitions of phage and their hosts from marine to terrestrial freshwater environments, without time for wider dispersal across continents. However, another factor could be sampling bias, as most of the samples analyzed were from North America (Fig.5).

The co-occurrence of HTVC010P-related phage and *Fonsibacter* suggests a stable host-phage relationship. However, despite the presence of *Fonsibacter* phage, we did not identify *Fonsibacter* in freshwater sediment or three out of the four hydraulically fracturing related samples (Fig.5, Supplementary Fig.11) or in one sample from the EPL water-sediment interface sample. The phage in the sediment samples could have settled from the overlying water column. As similar process may explain the presence of phage that shared identical TerL sequences in both the freshwater and sediment Lake Kivu samples (Supplementary Table 8).

### Multiple marine-freshwater transitions of *Fonsibacter* phage

The similarity between the *Fonsibacter* phage and Pelagiphage (Figs.1 and 2) indicates that they share a common ancestor. A question is whether *Fonsibacter* phage transitioned from marine environments only once or multiple times. We detected lineages 1 and 2a-e of the HTVC010P-related phage in many freshwater habitats and some environments with fresh water to saline gradients (Fig.5, Supplementary Table 8). However, the majority of lineage 3 of HTVC010P-related phage were from marine habitats, with only three TerL from two freshwater habitats (Fig.5). This observation, and their 99-100% similarity to those from marine/saline habitats, suggests a relatively recent transition of this lineage from marine/saline to freshwater ecosystems (Supplementary Information). On the other hand, detection of HTVC010P-related_33_76 at high relative abundance in EPL across an eight-months sampling period (Fig.3b, Supplementary Fig.6), indicates persistence in this habitat. Moreover, these four freshwater phage clustered into two distinct groups (Fig.5), possibly indicating that they originated from different marine phage genotypes. Similarly, given the global distribution of their marine relatives (Zhao et al. 2013; Zhao et al. 2018) versus relatively limited distribution of uv-Fonsiphage-EPL related phage in freshwater (Supplementary Fig.10) may point to their relatively recent transition from a marine to freshwater habitats.

## Conclusions

The persistent inertia in culturing freshwater microbes challenges our understanding of the ecology and functions of aquatic ecosystems. Genome-resolved metagenomics is a promising approach to solve this problem, by reconstructing complete genomes of bacterial hosts and their infecting phage. In this study, we report complete genomes of *Fonsibacter* and both lysogenic and lytic infecting phage, revealed their similarity to marine Pelagiphage and wide distribution in freshwater habitats. Based on this, more detailed analysis on the interaction of *Fonsibacter* and infecting phage could be performed in future studies.

## Methods

### Sampling, DNA extraction, sequencing, metagenomic assembly and genome binning

The EPL samples were collected from a end pit lake in Alberta of Canada in 2017 (15 samples) and 2018 (2 samples), at multiple depths (Supplementary Table 1). Also, one sample was collected from the input source of EPL (I-EPL) on September 19th, 2017 at a depth of 0.5 m (I-EPL_09192017_0.5m). The BMMRE samples were collected from a base metal mine receiving environment in northern Manitoba, Canada, in 2016 (July 24th) and 2017 (April 13th and September 27th). The geochemical features of the samples were determined *in situ* or in the laboratory as previously described (Risacher et al. 2018).

Genomic DNA was collected filtering ca. 1.5 L water through 0.22-µm Rapid-Flow sterile disposable filters (Thermo Fisher Scientific) and stored at −20 °C until the DNA extraction. DNA was extracted from the filters as previously described (Whaley-Martin et al. 2019). The DNA samples were purified for library construction, and sequenced on an Illumina HiSeq1500 platform with PE150 bp kits. The raw reads of each metagenomic sample were filtered to remove Illumina adapters, PhiX and other Illumina trace contaminants with BBTools (Bushnell 2018), and low-quality bases and reads using Sickle (version 1.33; https.github.com/najoshi/sickle). The high-quality reads of each sample were assembled using idba_ud (Peng et al. 2012) (parameters: --mink 20 --maxk 140 --step 20 --pre_correction). For a given sample, the high-quality reads of all samples from the same sampling site were individually mapped to the assembled scaffold set of each sample using bowtie2 with default parameters (Langmead & Salzberg 2012). The coverage of scaffold was calculated as the total number of bases mapped to it divided by its length. Multiple coverage values were obtained for each scaffold to reflect the representation of that scaffold in the various samples. For each sample, scaffolds with a minimum length of 1.5 kbp were assigned to preliminary draft genome bins using MetaBAT with default parameters (Kang et al. 2015), with both tetranucleotide frequencies (TNF) and coverage profile of scaffolds considered. The scaffolds from the obtained bins and the unbinned scaffolds with a minimum length of 1 kbp were uploaded to ggKbase (http://ggkbase.berkeley.edu/). The genome bins detected with *Fonsibacter-*related scaffolds were evaluated based on the consistency of GC content, coverage and taxonomic information and scaffolds identified as contaminants were removed.

### Manual curation of *Fonsibacter* and phage genomes

The *Fonsibacter* genome bin with 27 scaffolds was manually curated to completion, by firstly performing an overlap-based assembly of scaffolds using Geneious (Kearse et al. 2012), then linkage of scaffolds by metaSPades assembled scaffolds and scaffold extension, and manual fixation of local assembly errors detected by ra2.py (Brown et al. 2015). A total of 51 bacterial universal single-copy genes (SCGs) were used to evaluate genome completeness (Anantharaman et al. 2016). One prophage was detected in the complete *Fonsibacter* genome. This prophage was manually curated into a circular genome using paired-end reads located at both ends of the prophage region in the host genome. To obtain genomes of potential *Fonsibacter*-infecting phage, we identified the EPL/I-EPL/BMMRE scaffolds with multiple genes most close to those of published Pelagiphage. For these scaffolds, manual curation including assembly error fixation was performed (using the same methods as for the *Fonsibacter* genome). To investigate (and for reference) how the circular genome of the phage relate to the prophage sequence, see the step-by-step procedures in the Supplementary Information.

### Gene prediction, annotation, CRISPR-Cas analyses and protein families analyses

The protein-coding genes of the curated *Fonsibacter* and phage genomes were predicted using Prodigal (Hyatt et al. 2010), and searched against KEGG, UniRef100 and UniProt for annotation, and metabolic pathways were reconstructed. The 16S rRNA gene of *Fonsibacter* was predicted based on the HMM model as previously described (Brown et al. 2015). The tRNAs in *Fonsibacter* and phage genomes were predicted using tRNAscan-SE 2.0 (Lowe & Chan 2016). The transmembrane domains and single peptide of proteins were predicted using Phobius (Käll et al. 2007). The identification of CRISPR-Cas systems in assembled scaffolds was performed using python script (https://github.com/linxingchen/CRISPR), all unique CRISPR spacers were extracted from the scaffolds and reads mapped to the scaffolds, and searched against the curated phage genomes for the potential target using blastn (blastn-short). ANI was calculated using the online tool OrthoANIu (Lee et al. 2016).

For comparative genomic analyses of phage related to uv-Fonsiphage-EPL, we included all the published *HTVC019Pvirus* pelagiphage as analyzed in Zhao et al. (Zhao et al. 2018). For comparative analyses of HTVC010P-related phage, we searched the TerL proteins against NCBI-nr using BLASTp and NCBI scaffolds/genomes having a hit with ≥ 70% similarity (few with ≥ 80% similarity) were retained for further analyses (Supplementary Table 2). The predicted proteins of the selected NCBI scaffolds/genomes were downloaded from NCBI, protein families analyses were performed as previously described (Meheust et al. 2018), including the proteins of newly constructed phage genomes. In detail, first, all-vs-all searches were performed using MMseqs2 (Steinegger & Söding 2017), with parameters set as e-value = 0.001, sensitivity = 7.5 and cover = 0.5. Second, a sequence similarity network was built based on the pairwise similarities, then the greedy set cover algorithm from MMseqs2 was performed to define protein subclusters (i.e., protein subfamilies). Third, in order to test for distant homology, we grouped subfamilies into protein families using an HMM-HMM comparison procedure as follows. The proteins of each subfamily with at least two protein members were aligned using the result2msa parameter of MMseqs2, and HMM profiles were built from the multiple sequence alignment using the HHpred suite (Söding et al. 2005). The subfamilies were then compared to each other using hhblits (Remmert et al. 2011) from the HHpred suite (with parameters -v 0 -p 50 -z 4 -Z 32000 -B 0 -b 0). For subfamilies with probability scores of ≥ 95% and coverage ≥ 0.5, a similarity score (probability ⨉ coverage) was used as the weights of the input network in the final clustering using the Markov CLustering algorithm (Enright et al. 2002), with 2.0 as the inflation parameter. Finally, the resulting clusters were defined as protein families.

### Phylogenetic analyses

Multiple phylogenetic trees based on different gene (or gene sets) were built in this study (some are described in the Supplementary Information).

1. 16 ribosomal proteins (rps) of SAR11 genomes: for reference, SAR11 genomes at NCBI were downloaded and evaluated using CheckM to filter those genomes with completeness lower than 70%. The 16 rps (i.e., L2, L3, L4, L5, L6, L14, L15, L16, L18, L22, L24, S3, S8, S10, S17 and S19) were predicted from the NCBI genomes and the *Fonsibacter* genomes from this study, using HMM-based search as previously described (Anantharaman et al. 2016). Those genomes with none (AAA024-N17, AAA023-L09, AAA027-L15) or only two (AAA280-B11) of these 16 RPs were excluded for analyses.
2. ribosomal protein S3 (rpS3): this marker gene was used to identify *Fonsibacter* in metagenomic datasets, and also for phylogenetic analyses using the nucleotide sequences.
3. concatenated proteins of phage: via protein families analyses (see above), the 12 core proteins detected in the 28 uv-Fonsiphage-EPL related phage were used for phylogenetic analyses (Supplementary Table 3; two genomes lack one of the 12 core proteins due to incompleteness) (Zhao et al. 2018)).
4. TerL: the phage large terminase was used for several phylogenetic analyses, including the HTVC010P-related phage analyses, the phage detected in Lake Mendota (clustered with 99% identity) and the phage identified in other habitats worldwide.

For tree construction, protein sequences datasets were aligned using Muscle (Edgar 2004). All the alignments were filtered using TrimAL (Capella-Gutiérrez et al. 2009) to remove those columns comprising more than 95% gaps, and also ambiguously aligned C and N termini. For the 16 ribosomal proteins and the 12 phage proteins sets, sequences were concatenated into a single aligned sequence. The phylogenetic trees (including concatenated and TerL) were constructed using RAxML version 8.0.26 with the following options: -m PROTGAMMALG -c 4 -e 0.001 -# 100 -f a (Stamatakis 2015). For rpS3, the nucleotide sequences were aligned and filtered as described above, and the tree was built using RAxML version 8.0.26 with the following options: -m GTRGAMMAI -c 4 -e 0.001 -# 100 -f a (Stamatakis 2015). All the trees were uploaded to iTOL v3 for visualization and formatting (Letunic & Bork 2007).

### Codon usage analyses and co-occurrence of Fonsibacter and phage

The codon usage frequency of phage, bacterial and archaeal genomes was determined using cusp (Create a codon usage table) program of EMBOSS (The European Molecular Biology Open Software Suite), with protein-coding genes predicted by Prodigal (-m single, translation table 11). The prophage region in *Fonsibacter*_30_26 was removed from the host genome before performing gene prediction. Clustering analyses of all these genomes based on their codon usage frequency were performed using the R package of “pheatmap” (Kolde 2012), with “Euclidean” clustering and “average” method (Fig.3a). The usage frequency of each synonymous codon for a given amino acid is listed in Supplementary Table 5. To evaluate the occurrence of *Fonsibacter* and phage in the EPL/I-EPL/BMMRE samples (Fig.3b), we used the rpS3 gene to identify *Fonsibacter* (and also the three genomes had similar codon usage frequency), and the TerL gene to identify phage. The genotype was determined based on sharing ≥ 99% phage TerL or *Fonsibacter* rpS3 amino acid similarity.

### Analyses of published data from Lake Mendota

*Fonsibacter* was studied previously in Lake Mendota (Zaremba-Niedzwiedzka et al. 2013; Garcia et al. 2018). The published metagenomic datasets deposited at IMG were searched for phage similar to the ones from EPL/I-EPL/BMMRE samples, using their TerL proteins as queries. We also obtained the rpS3 protein sequences from these datasets using BLASTp at IMG and HMM-based confirmation using the TIGRFAM database (Haft et al. 2003). The rpS3 proteins belonging to *Fonsibacter* were identified by phylogenetic analyses with all available SAR11 rpS3 sequences.

Raw paired-end reads of the time-series Lake Mendota samples, were downloaded from NCBI SRA via the information provided in Garcia et al. (Garcia et al. 2018). A total of 90 datasets were available for download. Quality control was performed on those raw reads as described above. To determine the relative abundance of both *Fonsibacter* and phage in each sample, we first mapped quality-reads of each sample to all confirmed and non-redundant (clustered at 100% identity) rpS3 genes from the 90 Lake Mendota samples, then filtered the mapping file to allow no more than 3 mismatches for each read (equal to 98% similarity). The coverage of each rpS3 gene across all 90 samples was determined as described above. For a given sample, the total relative abundance of *Fonsibacter* was determined as the total *Fonsibacter* rpS3 gene coverage divided by the total rpS3 gene coverage. To have a similar evaluation of the abundance of phage in the samples, we did the mapping, filtering and coverage calculation for the non-redundant (clustered at 100% identity) TerL genes detected in all 90 Lake Mendota samples, using the same methods as used for *Fonsibacter* rpS3. In a given sample, the total relative abundance of *Fonsibacter* phage was calculated as the total coverage of TerL divided by the total coverage of all rpS3 genes.

We performed *de novo* assembly using idba_ud on the datasets from October 2012, in which the HTVC010P-related phage had a high coverage. BLAST (including BLASTp and BLASTn) was used to retrieve similar scaffolds to HTVC010P-related phage from the assembled datasets, followed by manual curation and resulting with one complete phage genome. Gene prediction and annotation were conducted as described above.

### Global search of similar phage in IMG metagenomic datasets

With the TerL proteins of uv-Fonsiphage-EPL and HTVC010P-related phage reported in this study as queries, a global search was performed against the metagenomic datasets in the IMG system using BLASTp. The hits were filtered with a minimum blast alignment coverage of 80% and a minimum similarity of 80%. We also searched for *Fonsibacter* in the metagenomic datasets with phage TerL detected, using the rpS3 protein sequences from all available *Fonsibacter* genomes as queries. The resulting hits were filtered ≥ 80% alignment coverage and ≥ 80% similarity (preliminarily determined that a given rpS3 with similarity < 85% to *Fonsibacter* rpS3 is not a *Fonsibacter* rpS3), then a phylogenetic tree was built to retrieve the sequences assigned to the *Fonsibacter* subclade. To report the BLASTp hits in the IMG metagenomic datasets in this manuscript, we asked the Principal Investigators for each public dataset for their permission to report the results (Supplementary Table 8). The data in these metagenomic datasets have been published (Denef et al. 2016; Otten et al. 2016; Pinto et al. 2016; Colatriano et al. 2018; Dalcin Martins et al. 2018; Tran et al. 2018; Daly et al. 2019), or are in preparation for publication (Davenport et al., in prep; Paver et al., in revision; Evans et al., in prep).

To compare the relationship of the related phage identified in this study and from IMG freshwater habitats, phylogenetic analyses based on TerL were performed. We also included the TerL detected as similarity outliers and those related to HTVC010P-related_33_76 from IMG marine/saline habitats. The TerL sequences were dereplicated from each sampling site using cd-hit (-c 1 -aS 1 -aL 1 -G 1). Then the representatives and the TerL from HTVC010P-related phage genomes reported in this study, and also those of HTVC010P and related marine phage (as references), were aligned and filtered for tree building (see the “**Phylogenetic analyses**” section). Another tree was built for the TerL of uv-Fonsiphage-EPL and relatives, with the same procedure as described above.

To evaluate the phylogeny and diversity of *Fonsibacter* in the samples with TerL detected and analyzed, phylogenetic analyses based on the rpS3 nucleotide sequences of *Fonsibacter* were performed. We also included all the *Fonsibacter* rpS3 identified in EPL/I-EPL/BMMRE samples, with rpS3 from SAR11 marine and brackish subclades as references. All the sequences were aligned and filtered for tree building (see the “Phylogenetic analyses” section in Methods).

## Acknowledgements

We thank Raphaël Méheust for his help in protein families analyses and constructive suggestions. We thank Brian Hedlund, Carmen Amaro, David Walsh, Duane Moser, James Stegen, Jessica Jarett, Kelly Wrighton, Konstantinos Konstantinidis, Maureen Coleman, Michael Wilkins, Rick Cavicchioli, Robert M. McKay, Sean Crowe, Stuart Jones, Timothy Otten, Timothy Y. James, Vincent Denef, and Wes Swingley for their permission to use the phage large terminase and *Fonsibacter* rpS3 sequences from their metagenomic datasets deposited at IMG. The study was supported by the Genome Canada Large Scale Applied Research Program and Ontario Research Fund – Research Excellence grants to L.A.W., and NSERC Canada and Syncrude Canada (Grant No. CRDPJ 403361–10).

## Author contributions

L.X.C. designed the study. J.F.M., G.L.J. and T.C.N. collected and prepared the samples for sequencing. T.C.N and L.A.W. provided the DNA sequencing. L.X.C. performed the metagenomic assembly, genome binning, genome annotation, phylogenetic analyses, protein families analyses, codon usage analysis, and global search of related phage. L.X.C. and J.F.B. performed manual genome curation to completion, and L.X.C. detailed the step-by-step procedures for phage genome. L.X.C. and J.F.B. wrote the manuscript. All authors read and approved the final manuscript.

## Conflict of interest

The authors declare that they have no conflict of interest.

## Data availability

The *Fonsibacter* and phage genomes have been deposited at NCBI under BioProject xxx. The genomes can also be downloaded from https://ggkbase.berkeley.edu/SAR11_and_Phage/organisms. Please note that it is necessary to register for a ggKbase account by providing an email address before accessing or downloading the data.

## Supplementary information

### Supplementary Tables

A separate Excel file includes all the Supplementary Tables, and below are the legends for the tables:

**Supplementary Table 1**. Geochemical characteristics of samples collected from EPL, I-EPL and BMMRE.

**Supplementary Table 2**. Information of published genomes of marine SAR11 (Pelagibacter) infecting phages.

**Supplementary Table 3**. The distribution of protein families in the HTVC019Pvirus phage genome.

**Supplementary Table 4**. The distribution of protein families in each HTVC010P-related phage genome.

**Supplementary Table 5**. Codon usage frequency of phage, bacterial and archaeal genomes reconstructed from EPL and I-EPL samples.

**Supplementary Table 6**. The detection of phages similar to those reported in this study in the published Lake Mendota data. The detected ones were sorted by sampling date and colored. Only those with a minimum identity of 90% are shown. These TerL were clustered based on 99% aa similarity using cd-hit before conducting the phylogenetic analyses, the representative sequences were indicated by “#”, the ones belonging to same cluster are in the same background color.

**Supplementary Table 7**. Coverage and relative abundance of Fonsibacter and phages in Mendota Lake.

**Supplementary Table 8**. General information of related TerL sequences obtained from IMG platform.

## Supplementary methods, results and discussion

### 1. The absence of ribosomal protein L30 is a shared feature of all published SAR11 genomes

When evaluated the completeness using the 51 single copy genes that are universal in bacterial genomes, we found the *Fonsibacter* genome reported in this study had no ribosomal protein L30 (rpL30) gene. Further analyses indicated all SAR11 but one (AAA795-11P) genome lacked the rpL30 gene. For reference, we retrieved the two rpL30 genes in the Alphaproteobacteria bacterium casp-alpha1, a megabin that most close to SAR11 (but not SAR11; (Mehrshad et al. 2016)). With these three rpL30 as queries, we searched NCBI using BLASTp for homologies. Phylogenetic analyses of the queries and their BLASTp hits indicated that, the one from AAA795-11P was clustered with those from Marinimicrobia, a candidatus group in the FCP superphylum (Supplementary Fig. 2a). Moreover, the other three genes on the scaffold of AAA795-11P with rpL30, were also most close to those from Marinimicrobia (Supplementary Fig. 2b). This result indicated that the rpL30 scaffold was misbinned into AAA795-11P. Thus, we concluded that the absence of rpL30 gene is a general feature of SAR11 genomes.

### 2. Alternatives for host cells lysis of SAR11 phages

No conventional lysozyme, key enzyme for an essential step in cell lysis of the virulence cycle, was detected for four phages in *HTVC019Pvirus* group I (including HTVC120P; Fig. 1c). However, a bacterial toxin which showed homology to phage lysozyme was present (Supplementary Table 3), and may perform this function (Patzer et al. 2012).

For HTVC010P-related phages, only two of them contained a lysozyme (Fig. 2b). Next to the lysozyme within these two genomes, we detected a holin (GTA_holin_3TM; PF11351) (Fig. 2a), which was also found in all other phage genomes excluding the FFC draft genome likely due to incompleteness (Fig. 2b). Holin is thought to enable lysis by providing access to the peptidoglycan (Hynes et al. 2012). In all but one of the phages without lysozyme, a gene encoding peptidase M15 (Peptidase_M15_3; PF08291) was detected next to the holin protein (Fig. 2b). Peptidase_M15_3 represents the C-terminal domain of zinc D-Ala-D-Ala carboxypeptidases from *Streptomyces* species (Charlier et al. 2011), and related peptidase with peptidoglycan hydrolase activity has been documented (Khakhum et al. 2016). Moreover, the prediction of these peptidase M15 using SWISS-MODEL indicated they matched with the 1lbu.1.A template, which was a muramoyl-pentapeptide carboxypeptidase for bacterial cell wall degradation. Based on this information, we speculated that in phage without lysozyme, the holin and peptidase M15 work together for lysis.

### 3. Potential reason for the low abundance of *Fonsibacter* phages in most Lake Mendota samples

Given the sampling strategy of Mendota Lake, that is, filtering microorganisms cells onto 0.2 μm pore-size filters (Garcia et al. 2018), and assuming that the HTVC010P-related phages had a comparative capsid size as the Pelagiphage HTVC010P (50 nm in diameter (Zhao et al. 2013)), we speculated that the obtained phages were primarily from the surface and/or inside the bacterial host cells. In this case, it is reasonable that the phages were with low relative abundance in most samples (Fig. 4c).

### 4. The sole example of CRISPR-Cas system in SAR11

To date, only one marine SAR11 genome (single-cell genome AAA240-E13) has been reported with a putative CRISPR locus (Thrash et al. 2014). However, no other cas protein was identified near the locus excepting a cas4-like protein located on another scaffold. We tried but failed to link these two scaffolds or with any other scaffold in the genome bin, based on sequence overlap at the scaffold ends. Given the wide detection of cas4-like genes in archaeal, bacterial and phage genomes (Hudaiberdiev et al. 2017; Chen et al. 2019), it remains unclear for the role of this only reported putative CRISPR-Cas system in SAR11.

### 5. Evidence for “the comparatively recently transition of lineage 3 of HTVC010P-related phages”

In the main text, we concluded that the lineage 3 of HTVC010P-related phages represented by HTVC010P-related_33_76 transitioned comparatively recently from marine to freshwater ecosystems (Fig. 5). Here we show study cases to support this conclusion.

#### 5.1 Case 1 - European eel may transition HTVC010P-related_33_76 and also marine SAR11

We detected one TerL in each of the two European eel related samples. The TerL were similar to that of HTVC010P-related_33_76 (80% and 82% amino acid similarity), and both of them were partial genes. To obtain full length TerL genes, we downloaded the raw reads from NCBI SRA under the accession number of SRR1586370 and SRR1586416 (Carda-Diéguez et al. 2014; Carda-Diéguez et al. 2015; Carda-Diéguez et al. 2017), which were sequenced with Illumina PE100 kit. Quality control was performed on the raw reads as described in the “Methods” section of the main text, followed by *de novo* assembly using idba_ud (parameters: --pre_correction --mink 20 --maxk 80 --step 20). The scaffolds were compared against the two incomplete TerL proteins using BLASTx, and the targeted scaffolds had complete TerL genes for both of them. The complete Terl shared high similarity with only one mismatch to that of HTVC010P-related_33_76 (Extended Data Fig. 1). We compared these two European eel related TerL proteins against all TerL proteins that we have identified in public databases, and found 12 of them had a minimum identity of 97% (up to 99.2%; Extended Data Fig. 2). These 12 TerL proteins were from Groves Creek Marsh (Skidaway Island, Georgia; sequences 3-8), White Oak River estuary (North Carolina, US; 9-13) and Delaware Coast (US; sequence 14).

It was hypothesized that the epidermal mucosa could work as a phage enrichment layer (Barr, Auro, et al. 2013; Barr, Youle, et al. 2013), and which was documented using the European eel (*Anguilla anguilla*) as an animal model (Carda-Diéguez et al. 2015; Carda-Diéguez et al. 2017). The European eel travel from Europe to the East Coast of North America and back to Europe during their life, they usually spawn and lay eggs in the Sargasso Sea (Aarestrup et al. 2009). Given the sampling site of these two European eel-related TerL proteins (sampled from Alfacada pond of Spain), and their high similarity to those detected in the East Coast of North America, we speculated that the European eel play a role in the transition of these phages into freshwater ecosystems in Europe. Moreover, the Sargasso Sea also plays a major role in the migration of the American eel and the American conger eel, we suspected if these eel species also have a similar role in the phage transition between the ocean and freshwater ecosystems in North America.

For the two European eel related samples, BLASTp search did not identify any proteins similar to *Fonsibacter* rpS3. We suspected if this is due to the low relative abundance of *Fonsibacter* in the corresponding communities. Upon this, we compared the quality-reads (see above) against all available SAR11 rpS3 nucleotide sequences with a minimum similarity of 80% using BLASTn (e-value threshold = 1e-10). As a result, no quality read showed the highest similarity to *Fonsibacter* rpS3 under these thresholds, we thus concluded that there was not *Fonsibacter* member in the sampled communities. However, we identified two identical rpS3 protein sequences assigned to the marine SAR11 subclade (scaffold_3267 and scaffold_1782; Extended Data Fig. 3). These proteins shared a 96% AA similarity to a Tara ocean protein (MGYP000052759201), via searching the Tara ocean database (https://www.ebi.ac.uk/metagenomics/sequence-search/search/phmmer)(no detailed geographic information of this sequence is available). Thus, we speculated that the European eel could also transition the marine SAR11 to freshwater ecosystems.

**Extended Data Fig. 1.**
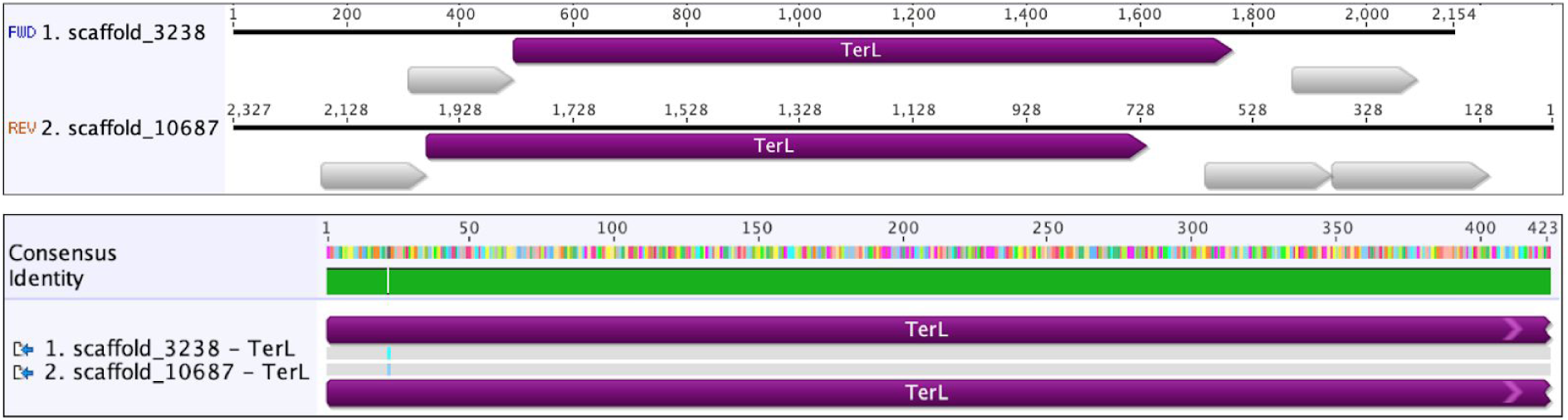
Upper panel: Genomic context of two European eel-related complete TerL (hypothetical proteins in gray). Bottom panel: Alignment of two European eel-related complete TerL proteins.

**Extended Data Fig. 2.**
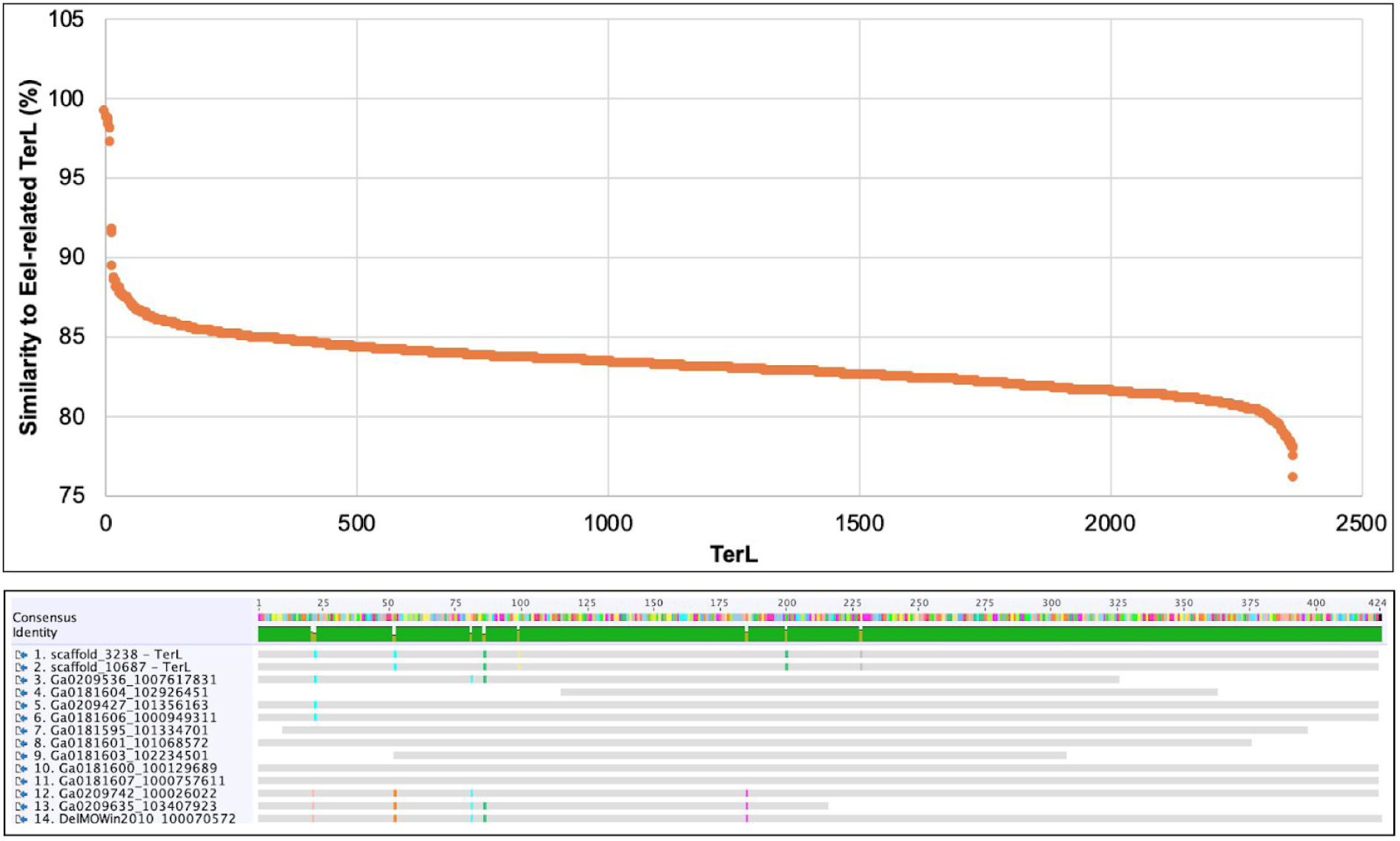
Upper panel: Similarity between European eel-related TerL proteins and those detected in global environments. Bottom panel: Alignment of European eel-related TerL proteins and relatives (> 97% similarity) from North America.

**Extended Data Fig. 3.**
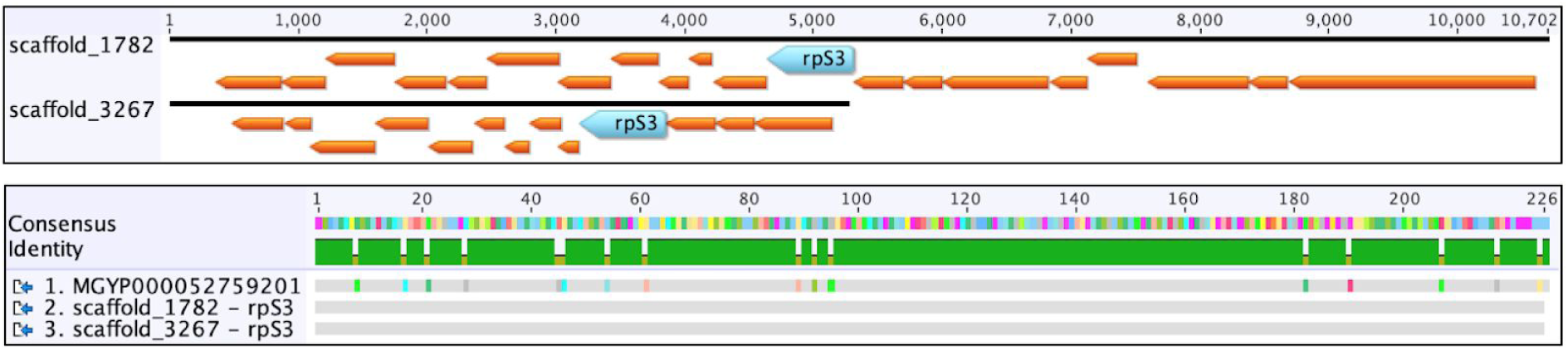
Upper panel: Genomic context of the SAR11 rpS3. Bottom panel: Alignment of the two identical SAR11 rpS3 proteins with one from Tara ocean project database.

#### 5.2 Case 2 - Identical HTVC010P-related_33_76 TerL found in a sediment sample of Lake Walker

The Lake Walker had a high concentration of total dissolved solids (TDS) due to the lower water level, which resulted from the overuse of water in the River Walker, the only input of Lake Walker (https://www.walkerbasin.org/history-of-walker-lake). As a result, the high TDS can no longer support the native fish and wildlife populations. For example, the Lahontan cutthroat trout have not been observed since year 2010 when TDS reached 20,000 mg/L. For reference, a TDS range of 8,000 - 12,000 mg/L is optimal for lake health conditions, typical seawater TDS is 40,000 mg/L, brackish TDS is around 10,000 mg/L.

One TerL from a Lake Warker sediment sample (collected on Nov 2nd of 2013; TDS = 19,000 mg/L) was identical to that of HTVC010P-related_33_76 (Extended Data Fig. 4a). We downloaded the raw reads from NCBI SRA (SRR5747948, generator of this unpublished data: Duane Moser), and performed quality control (as described in the “Methods” section of the main text). However, the relatively low abundance of this Lake Walker phage inhibited attempt for genome reconstruction, instead, we mapped the quality-reads to the HTVC010P-related_33_76 genome with 98% similarity. The result showed the high similarity of all the protein-coding genes with predicted function (Fig. 2), while not for some large genes encoding hypothetical protein (Regions 1 and 2; Extended Data Fig. 4b). By mapping EPL quality reads to the genome of HTVC010P-related_33_76 with 98% similarity, we detected similar coverage profiles as observed for the Lake Walker sample that, most of the EPL samples had a lower coverage in Regions 1 and 2. This observation likely indicated the occurrence of a close phage without the genes in Regions 1 and 2.

We had no clue when the Lake Walker phage started to appear. It is possible this phage transitioned to Lake Walker after the TDS has already raised too high to support lake fish and wildlife populations. In this case, it is more easy for this phage to adapt the habitat. Anyhow, the high similarity of Lake Walker reads (≥ 98%) mapped to genes of HTVC010P-related_33_76, indicated the Lake Walker phage is very close to HTVC010P-related_33_76.

**Extended Data Fig. 4.**
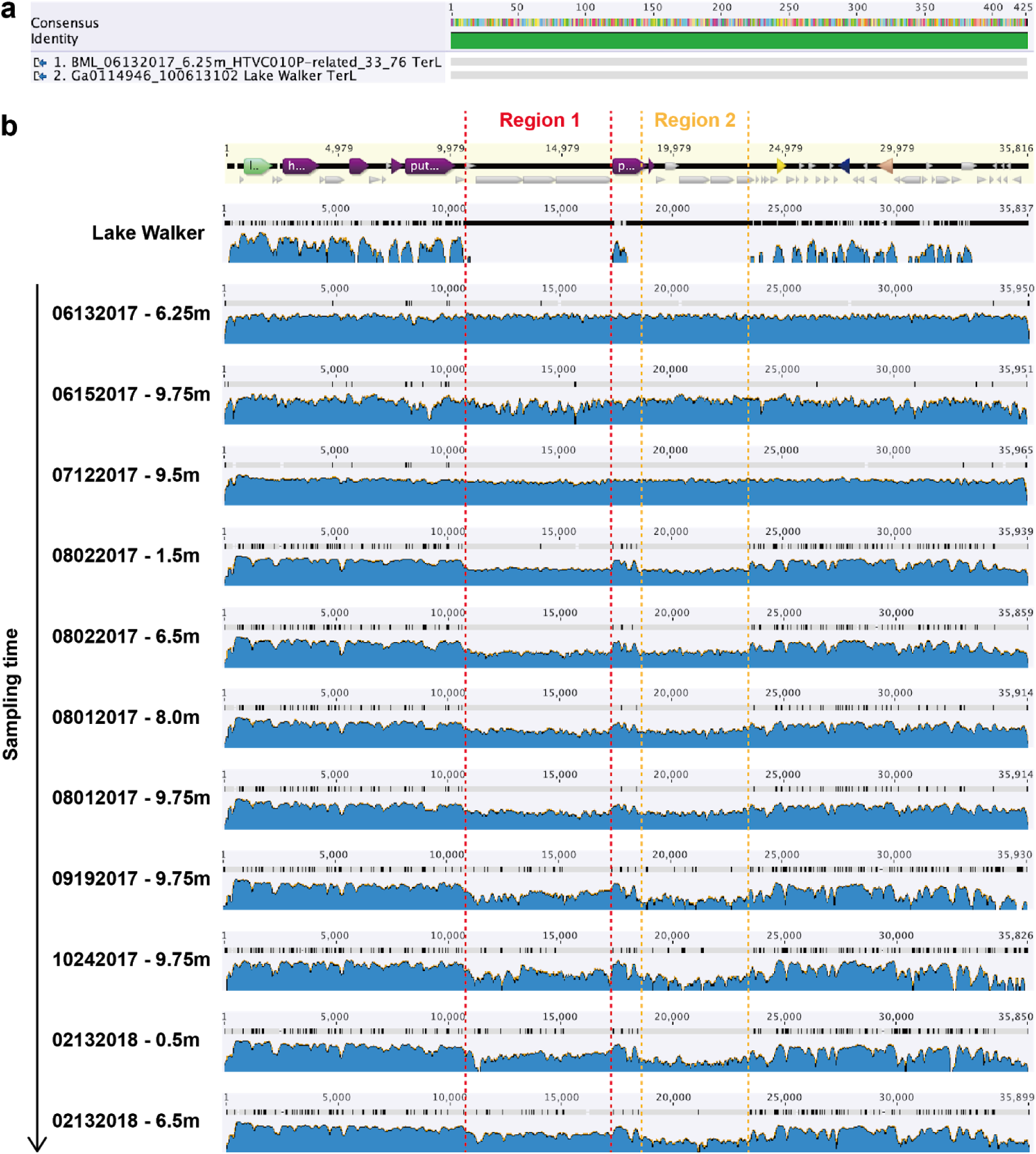
Comparative analyses related phages from EPL (HTVC010P-related_33_76) and Lake Walker sediment. (a) Identical TerL of HTVC010P-related_33_76 and the related phage from Lake Walker. (b) mapping of paired-end reads from the Lake Walker sediment sample, and different sampling points (and depth) to the genome of HTVC010P-related_33_76 (with 98% read similarity to the phage genome).

#### 6. A guide to obtaining complete phage genome from prophage-containing scaffold

Some phages could perform the lysogenic life strategy by inserting the genome into their bacterial hosts, which are prophages. Once we obtain a bacterial host scaffold with prophage, it generally indicates that most of this phage population perform lysogenic life strategy (otherwise the phage genome will be in a separated scaffold, not in the host scaffold). However, in the same community or other related communities (collected at different time points or different locations within the same region), may exist the same host cells without the prophage. Thus, we may be able to confirm the exact recombination site of phage genome into the host genome, and also the true length of the phage genome. Here we show a detailed guide of how to obtain complete phage genome from the prophage genome in the host.

Step 1: when phage-specific proteins were identified in a scaffold of a bacterial/archaeal genome bin, paired-end reads should be mapped to this scaffold (using bowtie or similar tools), to confirm if there is a complete prophage genome in this scaffold (it means the prophage genome is in the middle of the scaffold; bacterial/archaeal genes + phage genes + bacterial/archaeal genes).

Step 2: check the mapping profile to see if there are paired-end reads spanning the potential prophage region (Extended Data Fig. 5). If existed, these paired-end reads mapped on the host scaffold were from host cells without the prophage, and these are in the prophage were from lytic phage cells. In this example, there are only two paired-end reads in the prophage region.

**Extended Data Fig. 5.**
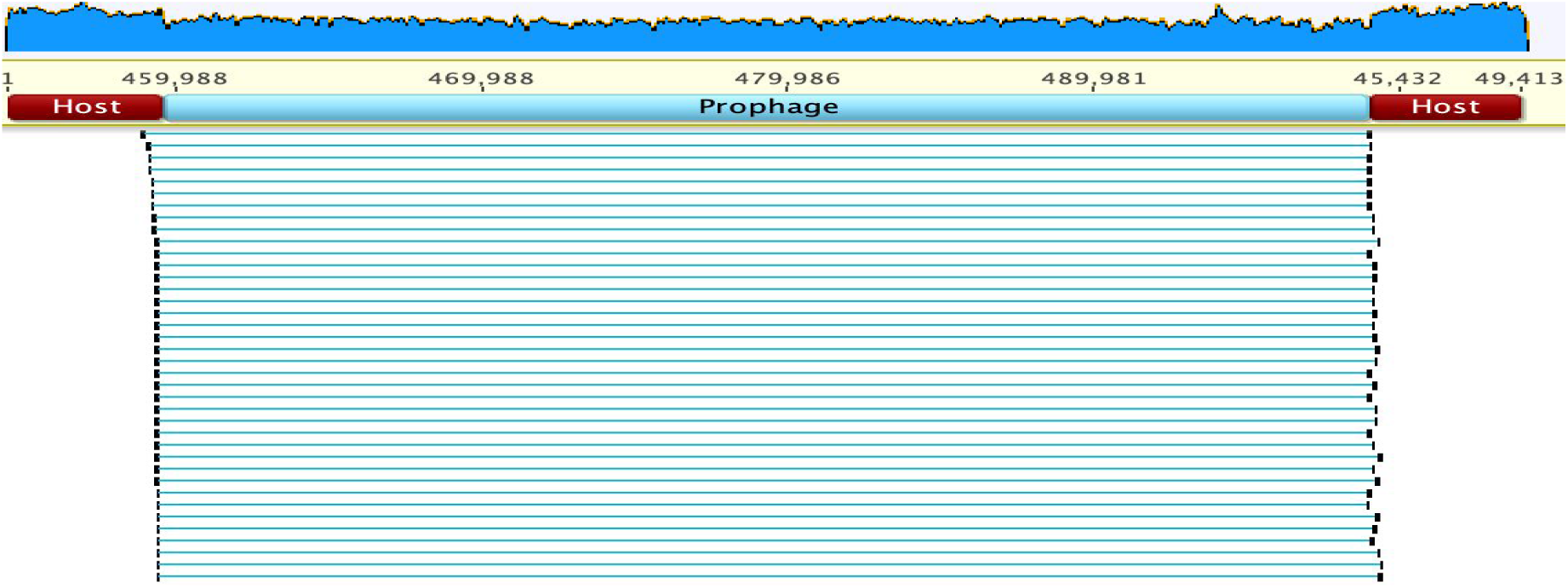
Long-distance spanning paired reads showing the location of prophage in host genome scaffold.

Step 3: the key step is to find some reads (multiple should be better) that with part of them perfectly matched to the scaffold, while the other part could not be matched, also the unmatched part from these partially-mapped reads could be aligned perfectly (Extended Data Fig. 6). In this example, “CTGGTATCGAAAGATGTGAAGGTTCAAGTCCTTTCTCCCGCACCATGTAGAATATGAAAGTA” (see the red zoom-in). Note that these reads were from the host cells without prophage. Upon this, the unmatched part must be found near the right end of the prophage on the scaffold (see the blue zoom-in). If this is true, then the recombination site could be determined as shown in Extended Data Fig. 6.

**Extended Data Fig. 6.**
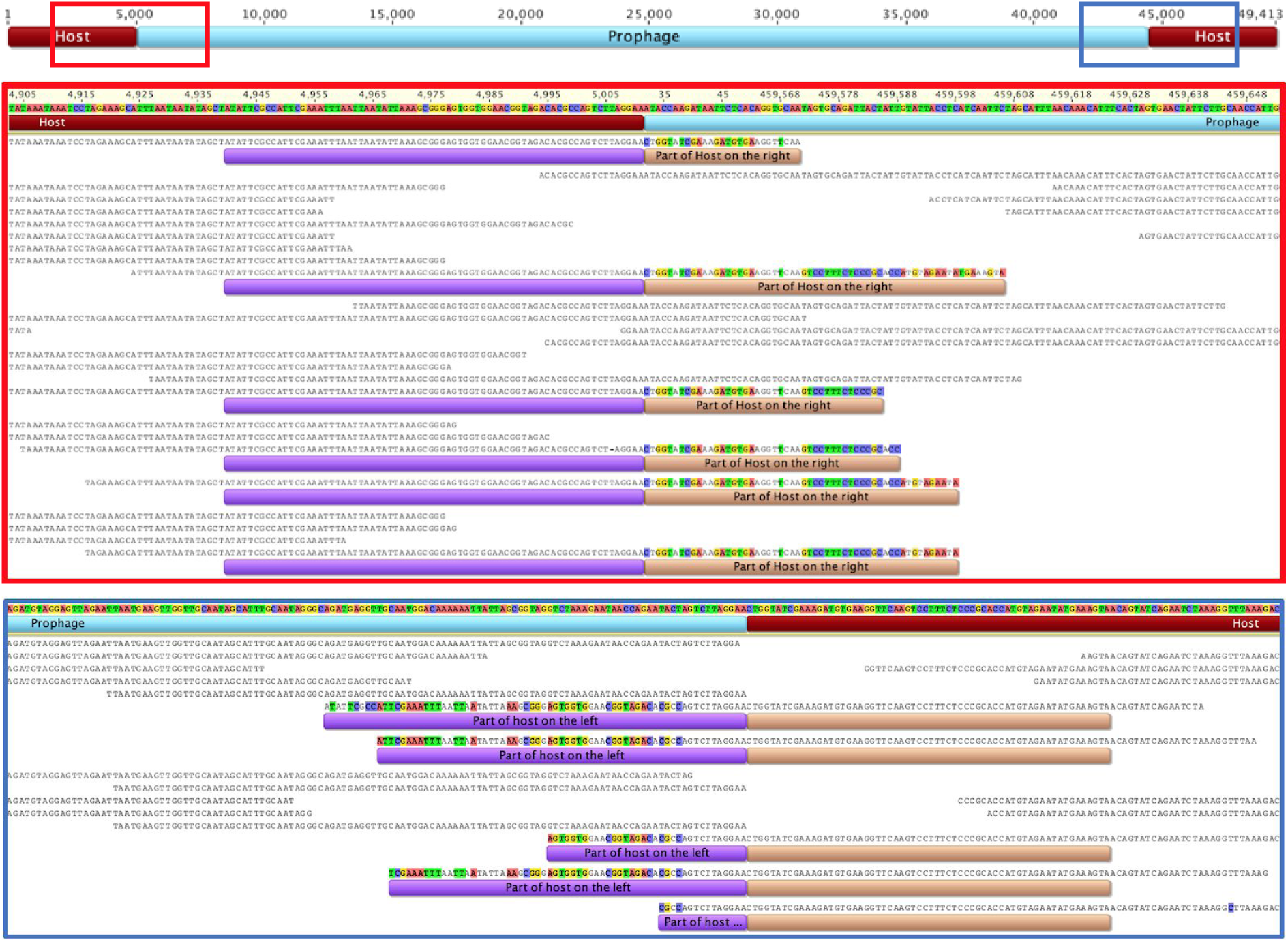
Upper panel: Overview of an example for a host scaffold with prophage. Middle panel: How to determine the left edge of prophage into host genome. Bottom panel: How to determine the right edge of prophage into host genome

Generally, we could do this by starting from the left end, or from the right end. However, there is a small difference if started from the right end, as we could see the unmatched part is not exactly at the end of the prophage (blue zoom-in box; Extended Data Fig. 6), this is because the phage shared a small part of DNA with the host (Extended Data Fig. 7), that is “AGTCTTAGGAA” (‘core sequence’), which is part of the host tRNA-Leu sequence, and also the recombination site. This recombination site sequence is with only one mismatch from that of Pelagiphage HTVC025P (“GTCTTAGGAAC”, (Zhao et al. 2018)).

**Extended Data Fig. 7.**
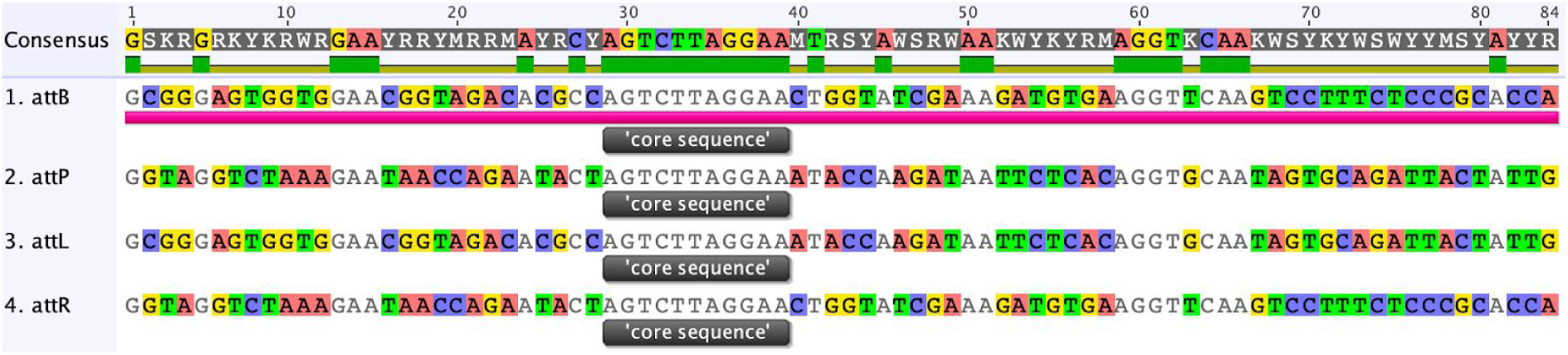
The ‘core sequence’ at the insertion site of prophage into host genome. The attB, aatP, attL and attR locations are shown and the recombination site was identified by sequence alignment.

Step 4: delete the prophage from the host genome scaffold based on the recombination site determined in step 3, and mapped quality reads to the host scaffold without prophage, to confirm the insertion location. Paired reads mapped to this region confirmed the insertion position (Extended Data Fig. 8).

**Extended Data Fig. 8.**
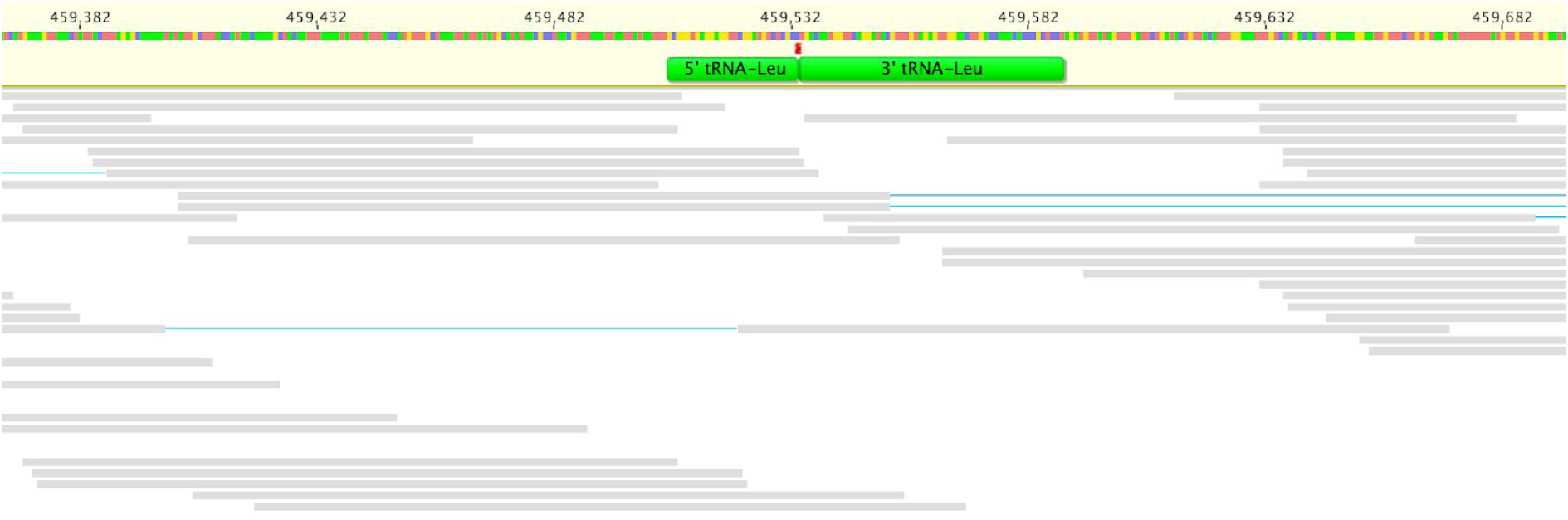
Reads mapping to the host scaffold with prophage deleted. The red stripe shows the recombination site of the phage into the host genome, which split the host tRNA-Leu into two fragments.

Step 5: map paired-end reads to the prophage genome to confirm if it is circular. If the prophage could be circularized, there should be reads that cover both end of the prophage genome. As shown below, we found three such reads (indicated in pink) (Extended Data Fig. 9). Also, we found two paired reads that flank both ends of the prophage genome. This information suggests the existence of free living particles of this phage.

**Extended Data Fig. 9.**
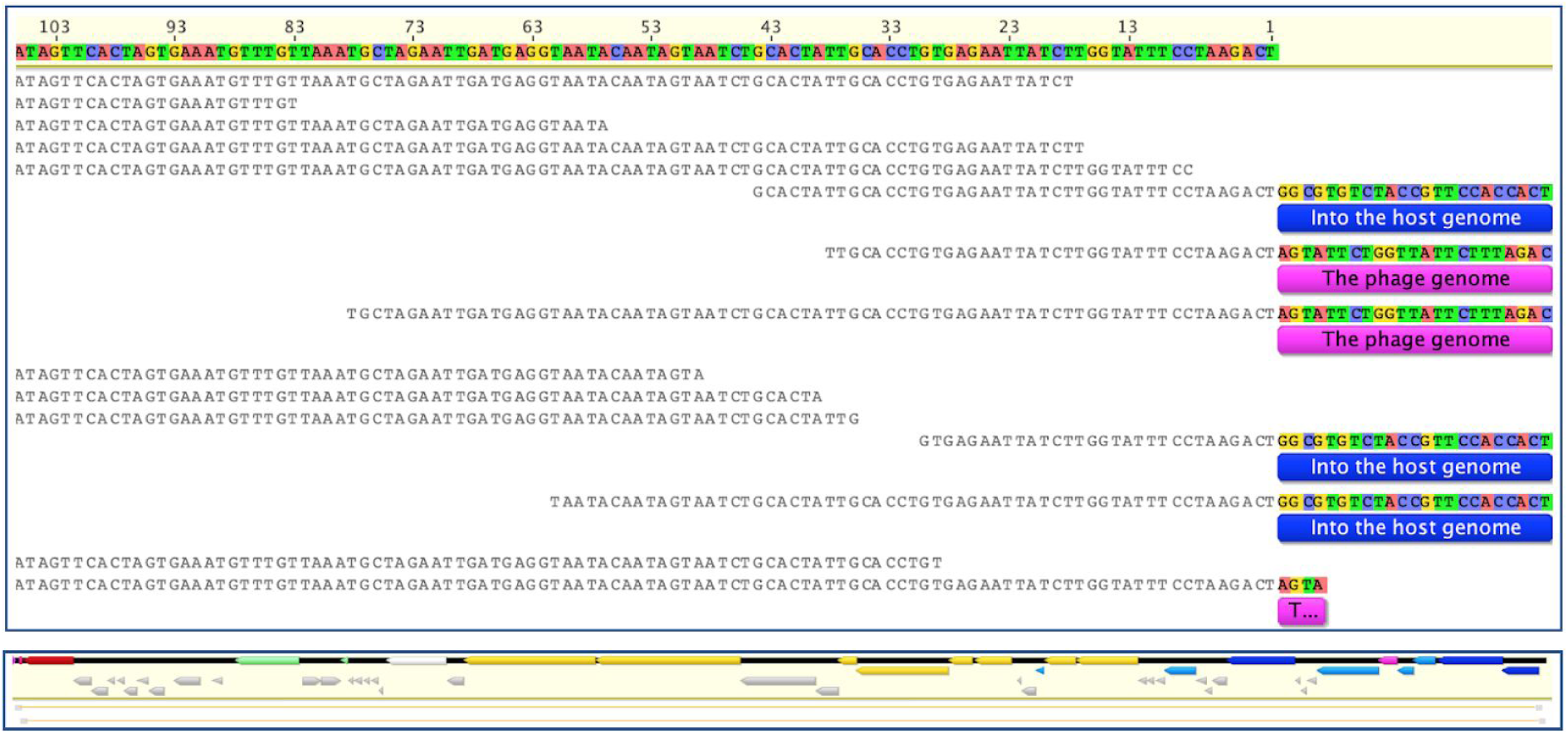
Upper panel: Reads mapping shows the prophage genome is circular. Bottom panel: Reads mapping shows end-to-end flanking paired reads.

It should be noted that under some circumstances the recombination site may not be determined, and/or the prophage could not be circularised into a complete phage genome, for example, (1) when all the host cells are with the prophage, (2) when the sequencing coverage is too low and there are not enough reads spanning the recombination sites, (3) when the prophage has several variants that share high sequence similarity, sometimes it will be impossible to determine.

## Supplementary Figures

**Supplementary Fig. 1.**
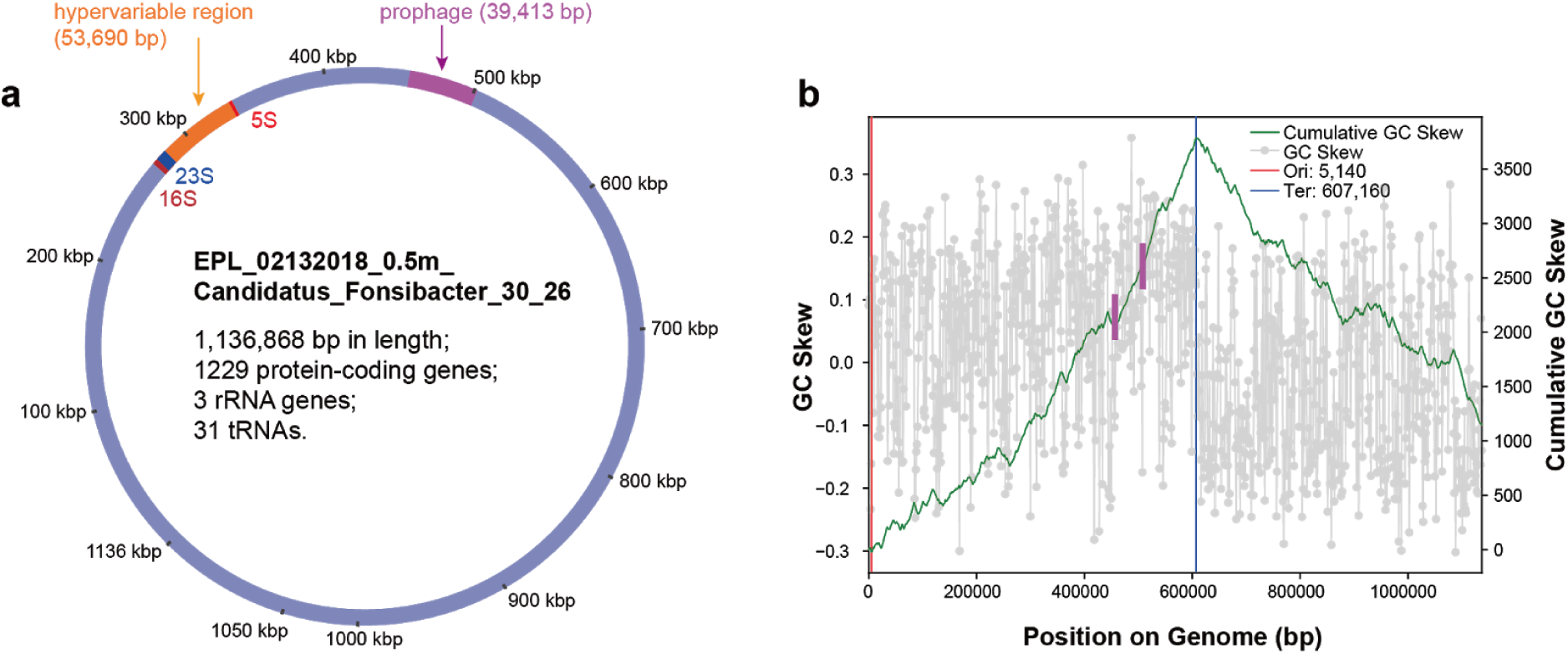
The complete genome of *Fonsibacter*_30_26 with a prophage. (a) The location of the prophage on the chromosome of its host. The location of 5S, and 23S/16S rRNA genes and the HVR2 are also shown. (b) The GC skew of the complete genome. The position of prophage is indicated by two purple lines.

**Supplementary Fig. 2.**
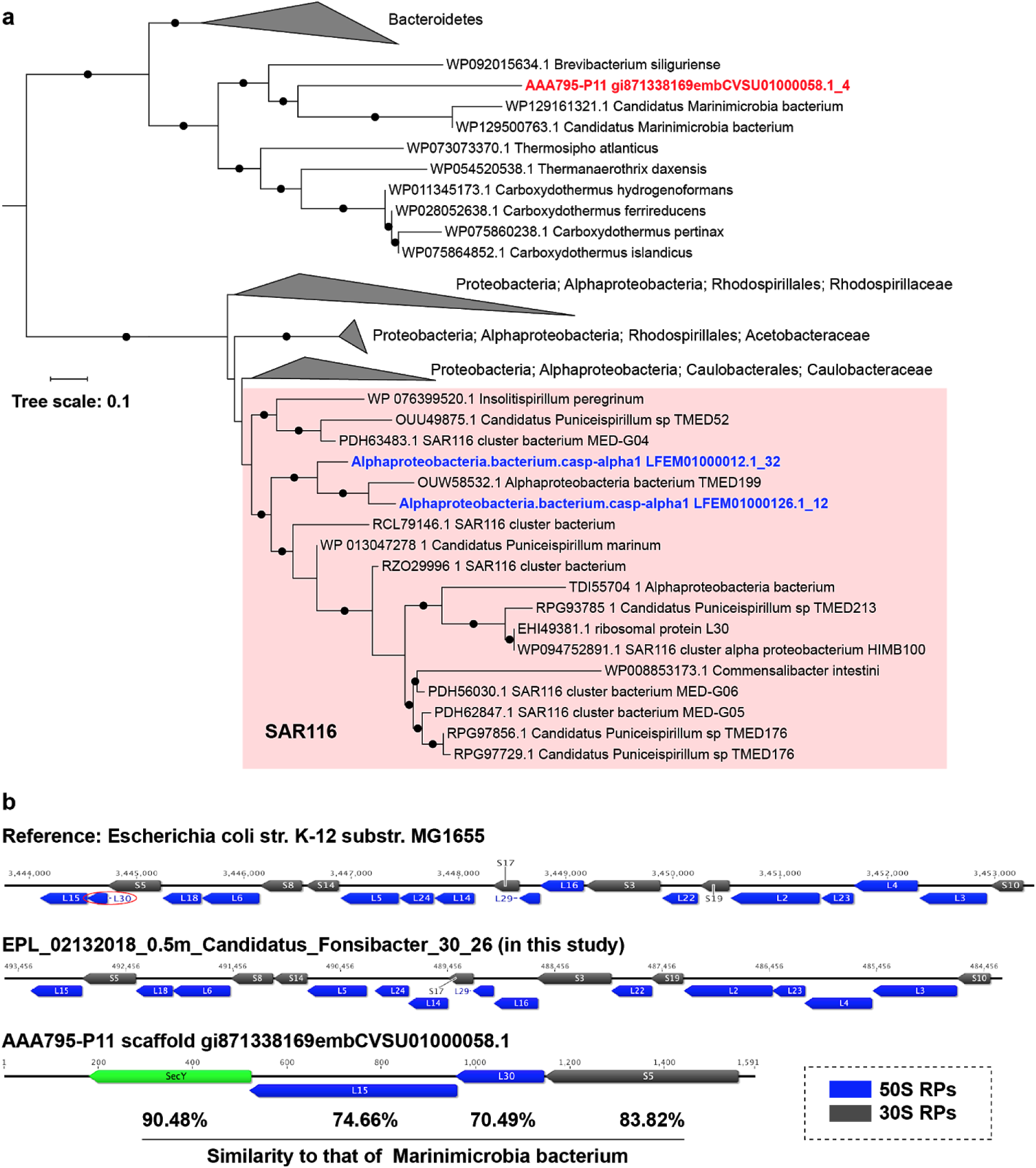
The absence of rpL30 gene in SAR11 genomes. (a) Phylogenetic analyses of the rpL30 identified in SAR11 genomes of AAA795-P11 (in red and bold). The rpL30 were determined based on TIGRFAMs HMM database search using HMMsearch within all published SAR11 genomes. The bootstraps value ≥ 70 is indicated by a black dot. (b) Detailed information of rpL30. The location of rpL30 on chromosome of E.coli (as reference; red circle). The absence of rpL30 in *Fonsibacter*_30_26 is shown in the middle. The scaffold with a rpL30 in AAA795-P11 is shown at the bottom. However, all four genes are most close to those from Marinimicrobia bacterium spp..

**Supplementary Fig. 3.**
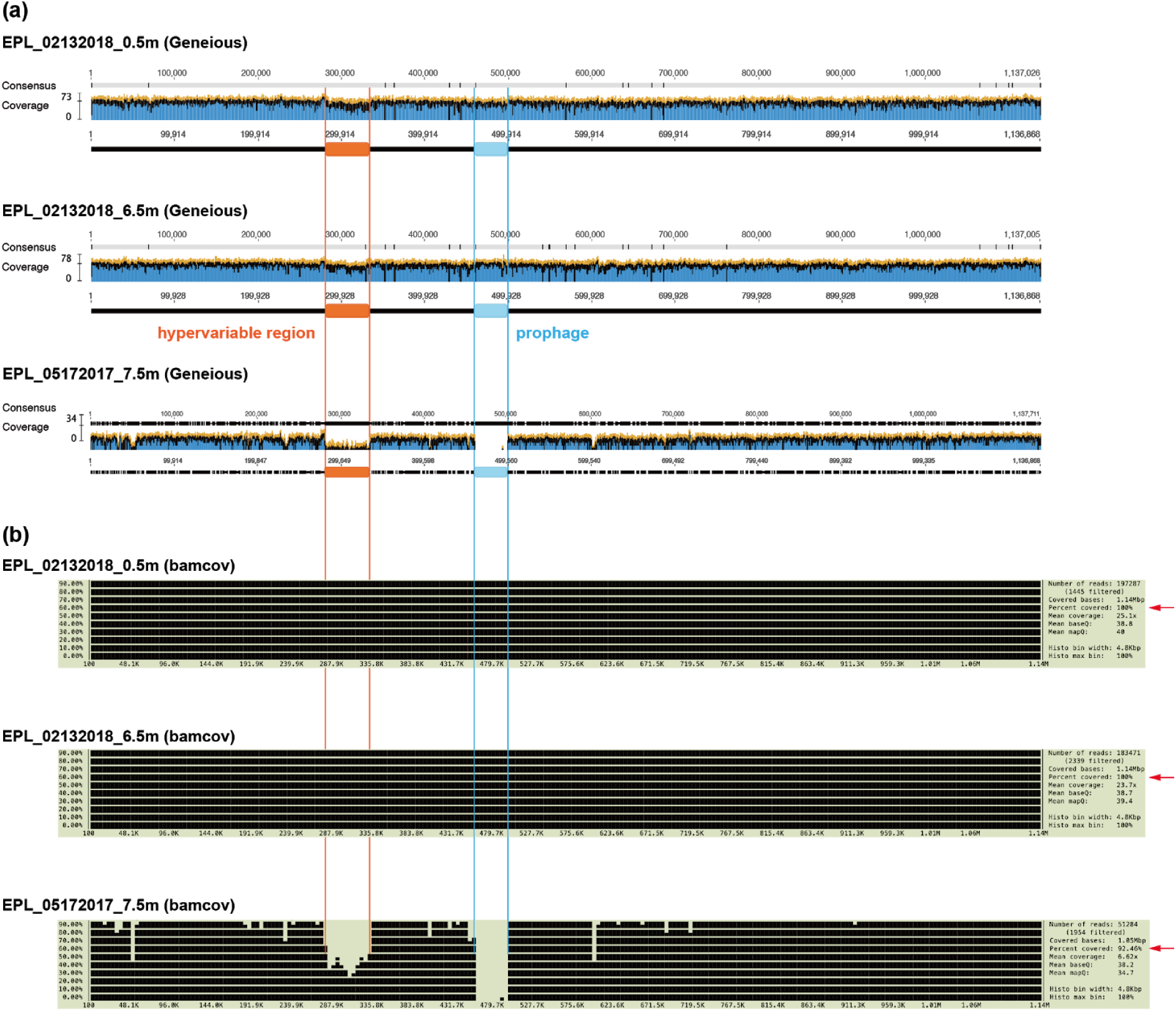
Mapping of paired-end reads from EPL samples to the complete genome of *Fonsibacter*_30_26, using (a) Geneious (Kearse et al. 2012) and (b) bamcov (https://github.com/fbreitwieser/bamcov). The hypervariable and prophage regions in the genome are highlighted. The hypervariable region was documented by mapped paired-end reads from the EPL_05172017_7.5m sample, and similar (pro)phage may be in this sample.

**Supplementary Fig. 4.**
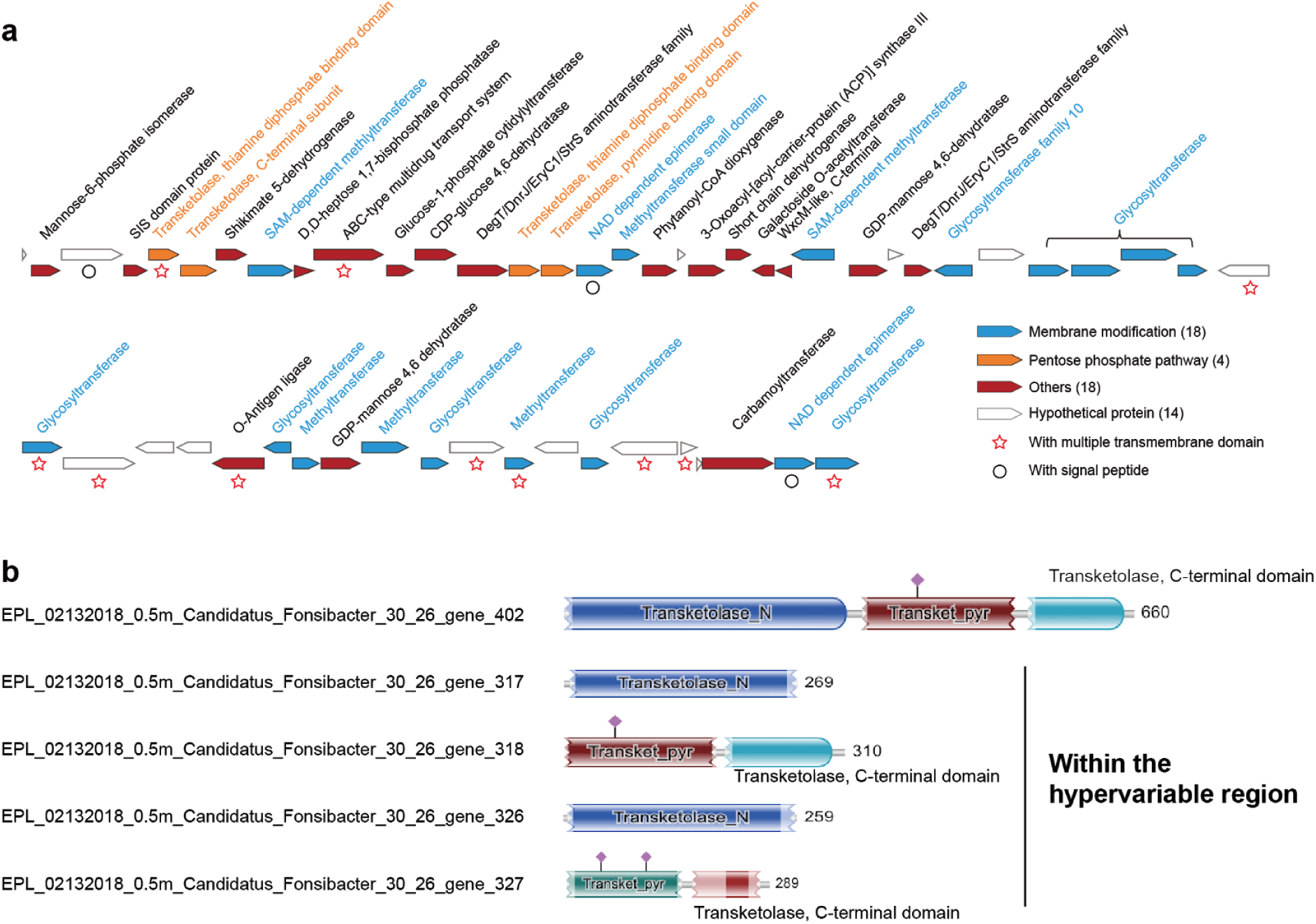
The hypervariable region of *Fonsibacter*_30_26. (a) Annotation of protein-coding genes in the hypervariable region. The genes were assigned to different functional categories and shown in different colors. The transmembrane domain and signal peptide were predicted and shown if identified, by a star and an open circle, respectively. (b) The four incomplete transketolase proteins in the hypervariable region. The full-length transketolase detected in the genome of *Fonsibacter*_30_26 but outside the hypervariable region is shown for comparison. Two of the transketolase proteins from the hypervariable region only have the N-terminal domain, and the other two have the transket_pyr and C-terminal domains.

**Supplementary Fig.5.**
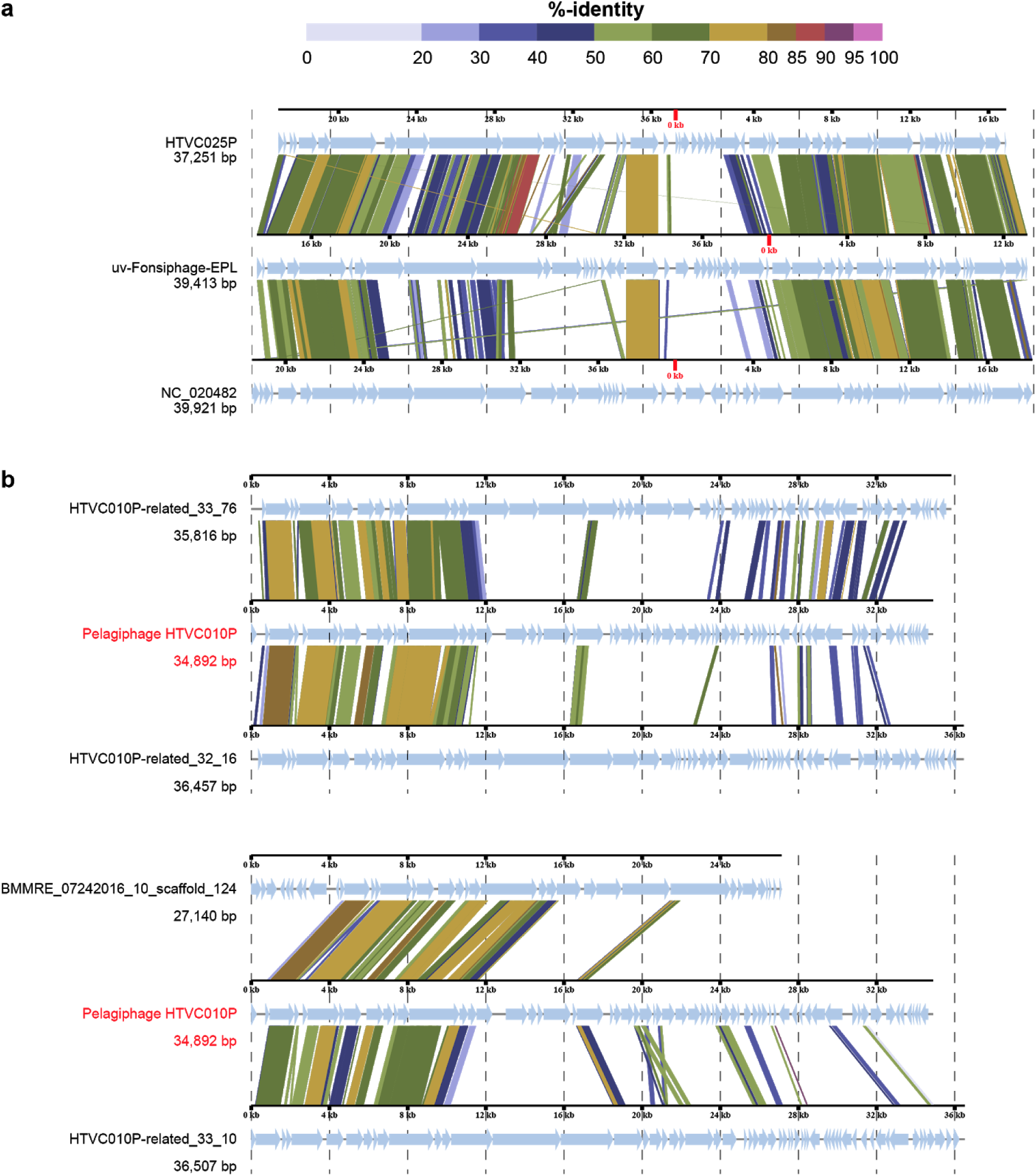
The alignment of phage genomes for (a) uv-Fonsiphage-EPL, HTVC025P and HTVC010P, and (b) HTVC010P and the three complete and one draft HTVC010P-related phage. The alignment was generated using viptree (https://www.genome.jp/viptree/) (Nishimura et al. 2017) with default parameters.

**Supplementary Fig. 6.**
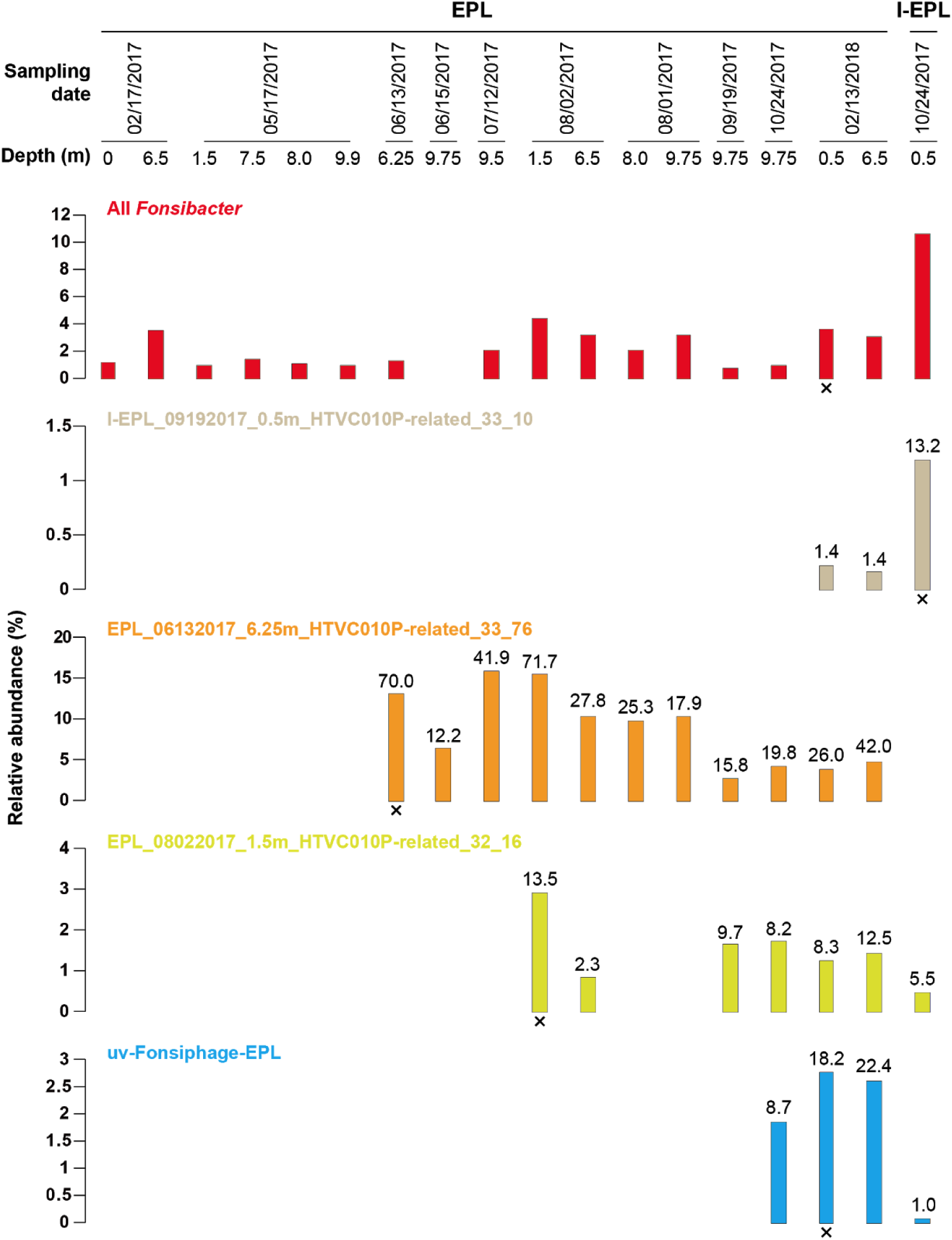
Relative abundance of *Fonsibacter* (accumulated) and infecting phages with complete genomes reconstructed from EPL and I-EPL samples. See Table 1 in the main text for information of complete genomes, the sample from where the complete genome was reconstructed is indicated by a “X”. The sequencing coverage of the phage genome is given above the bar. The calculation of relative abundance was performed as did for the Lake Mendota data (see Methods in the main text).

**Supplementary Fig. 7.**
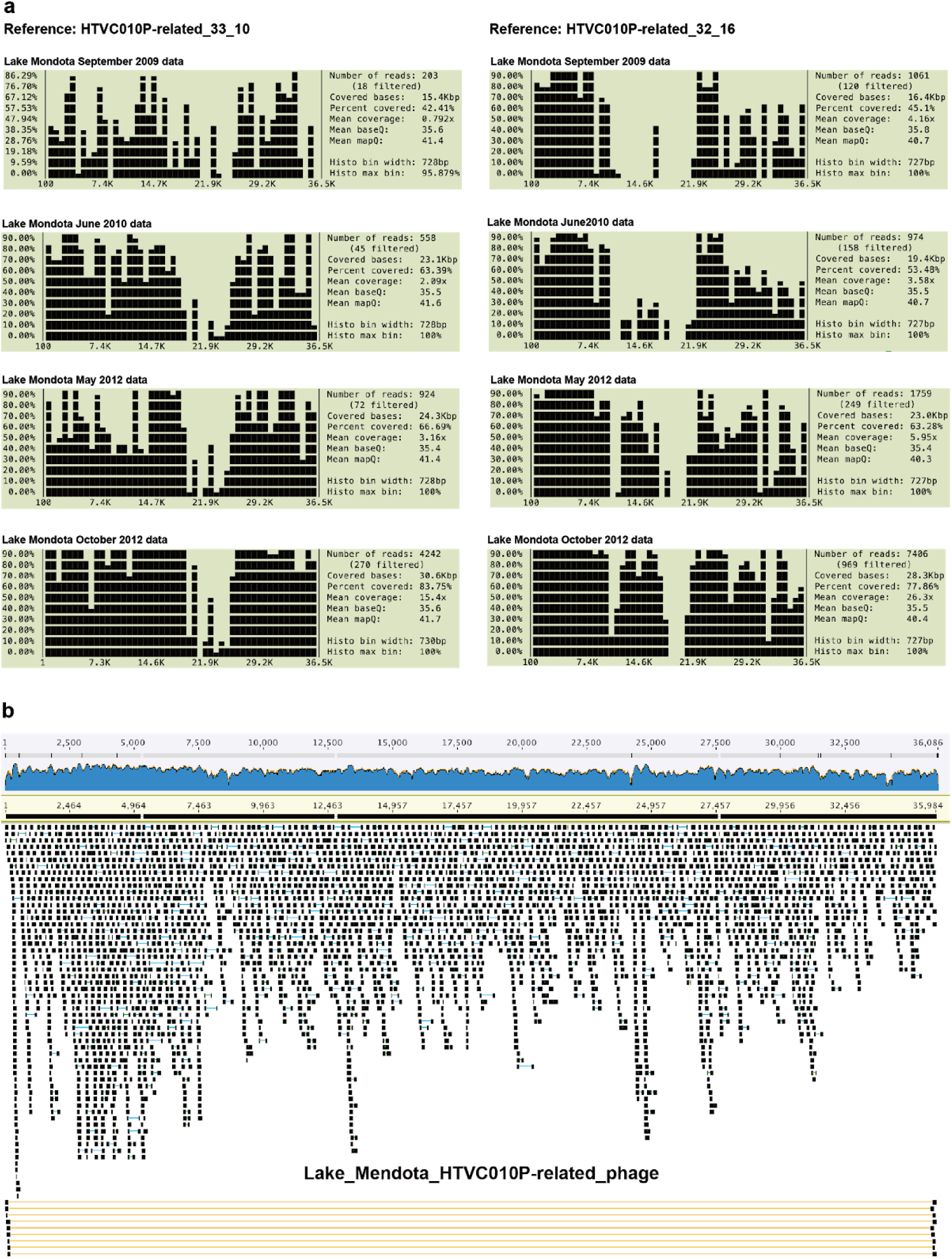
Related phages in Lake Mendota samples. (a) Mapping of paired-end reads from Lake Mendota samples to genomes of phages reported in this study. The mapped reads have been filtered with a minimum similarity of 98% to the reference genomes. Check Supplementary Table 5 for more details of the samples from Lake Mendota. (b) The reconstruction of a complete phage highly similar to EPL_08022017_1.5m_HTVC010P-related_32_16. Paired-end reads spanning the ends of the scaffolds are shown, and none local assembly error or Ns (gap) was reported by ra2.py, suggesting the 100% completeness of this phage genome.

**Supplementary Fig. 8.**
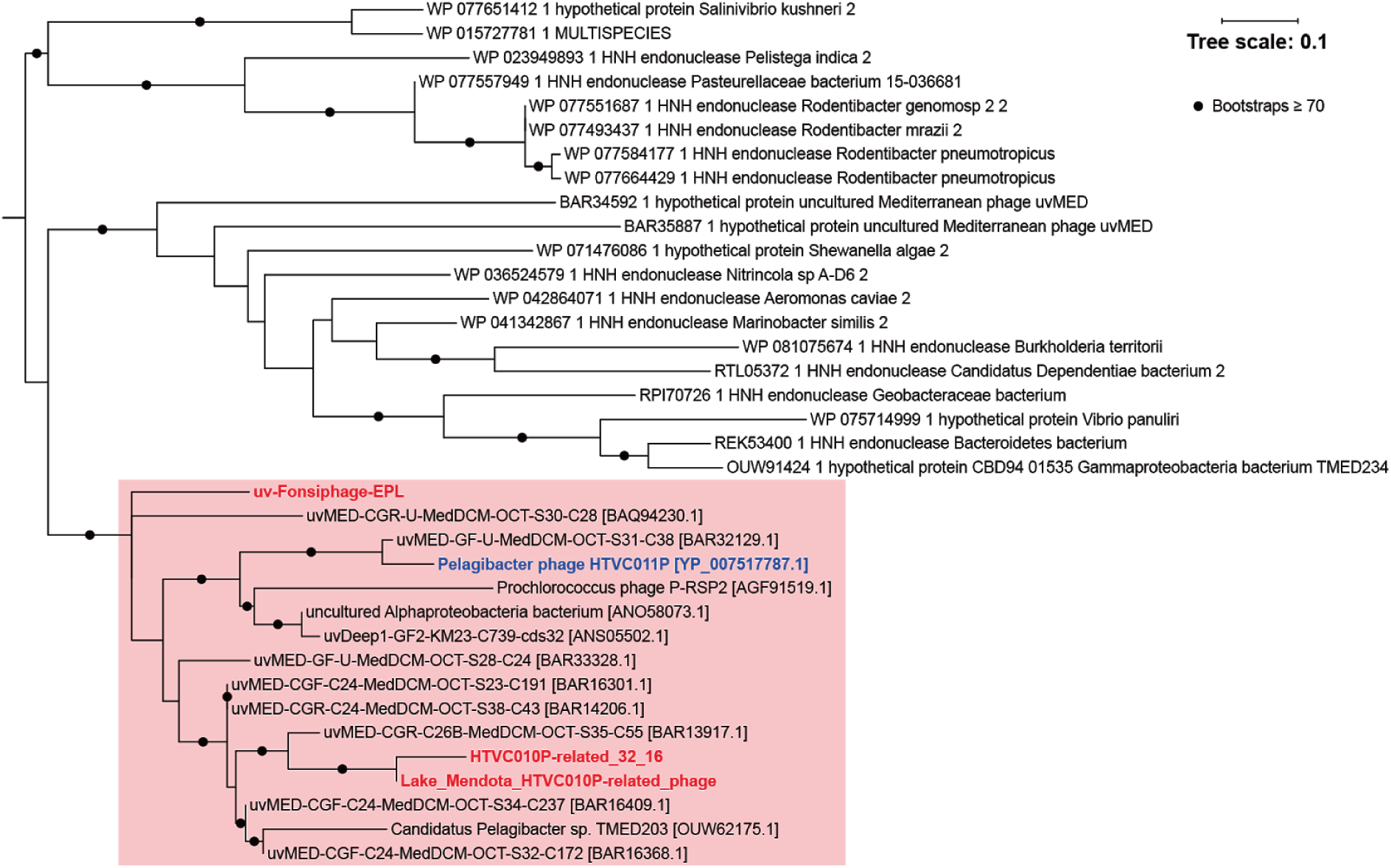
Phylogenetic analyses of HNH endonuclease identified in *Fonsibacter* phages. The HNH endonucleases of EPL_32_16 and Lake_Mendota were as queries for NCBI BLASTp search, the hits with >= 80 alignments and >= 50% similarity were retained for analyses, with the one from the prophage included as well. Note that one of the HNH endonucleases was from Candidatus Pelagibacter sp. TMED203 (Tully et al. 2017), the corresponding scaffold ID is NHJA01000005 at NCBI (https://www.ncbi.nlm.nih.gov/nuccore/NHJA01000005.1/). This scaffold contained many genes encoding hypothetical proteins and had a peptidase M15 gene, which was identified in most HTVC010P-related phages and thought to perform host cell lysis (see above). We suspect this is a misbinning of a phage scaffold into the genome bin of its bacterial host, or this scaffold represented a fragment of prophage in the host genome. We tried to assemble all the scaffolds (46 in total) in the genome bin TMED203 but failed to link NHJA01000005 to any of the others.

**Supplementary Fig. 9.**
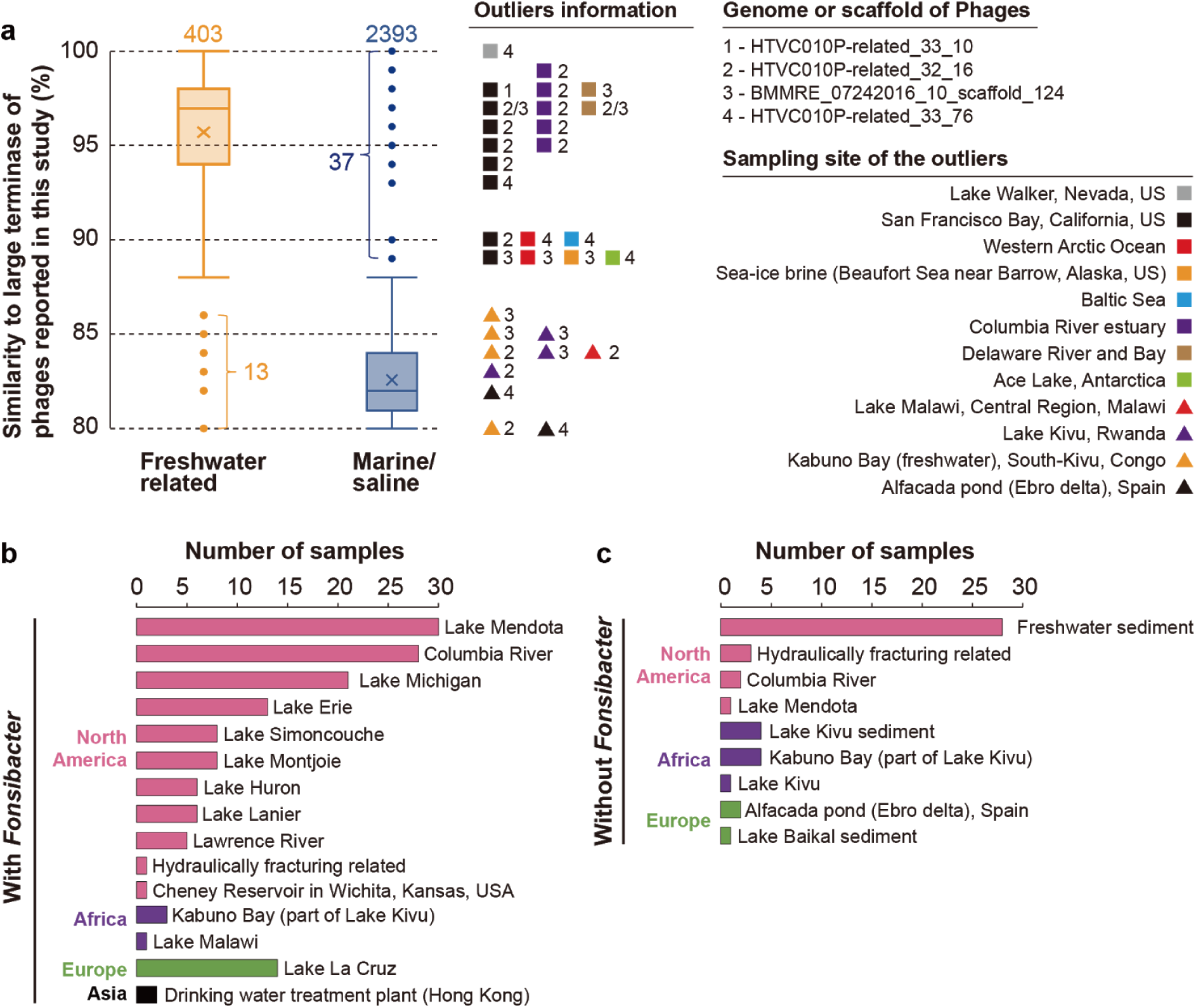
The detection of phages related to the ones reported in this study in global freshwater and marine/saline habitats. This analysis was performed by BLASTp search at IMG metagenomic datasets, with the TerL of phages reported in this study as queries, and only those with a minimum amino acid similarity of 80% were retained for analyses. (a) Box plots show the similarity of TerL detected in IMG datasets to those reported in this study, the numbers of total detections are shown above the boxes. For the outliers, their sampling sites (colored triangles for freshwater, and colored squares for marine/saline) and related phages (indexed number) are shown on the right. (b) Cooccurence of phage (based on TerL) and *Fonsibacter* (based on rpS3) in the sample. For each sampling site, the total number of samples with phage-*Fonsibacter* cooccurence were summed and shown. Sampling sites were grouped and colored by continental plates. (c) Samples with phage TerL detected while without *Fonsibacter* in the corresponding sample. Data was prepared as described in (b).

**Supplementary Fig. 10.**
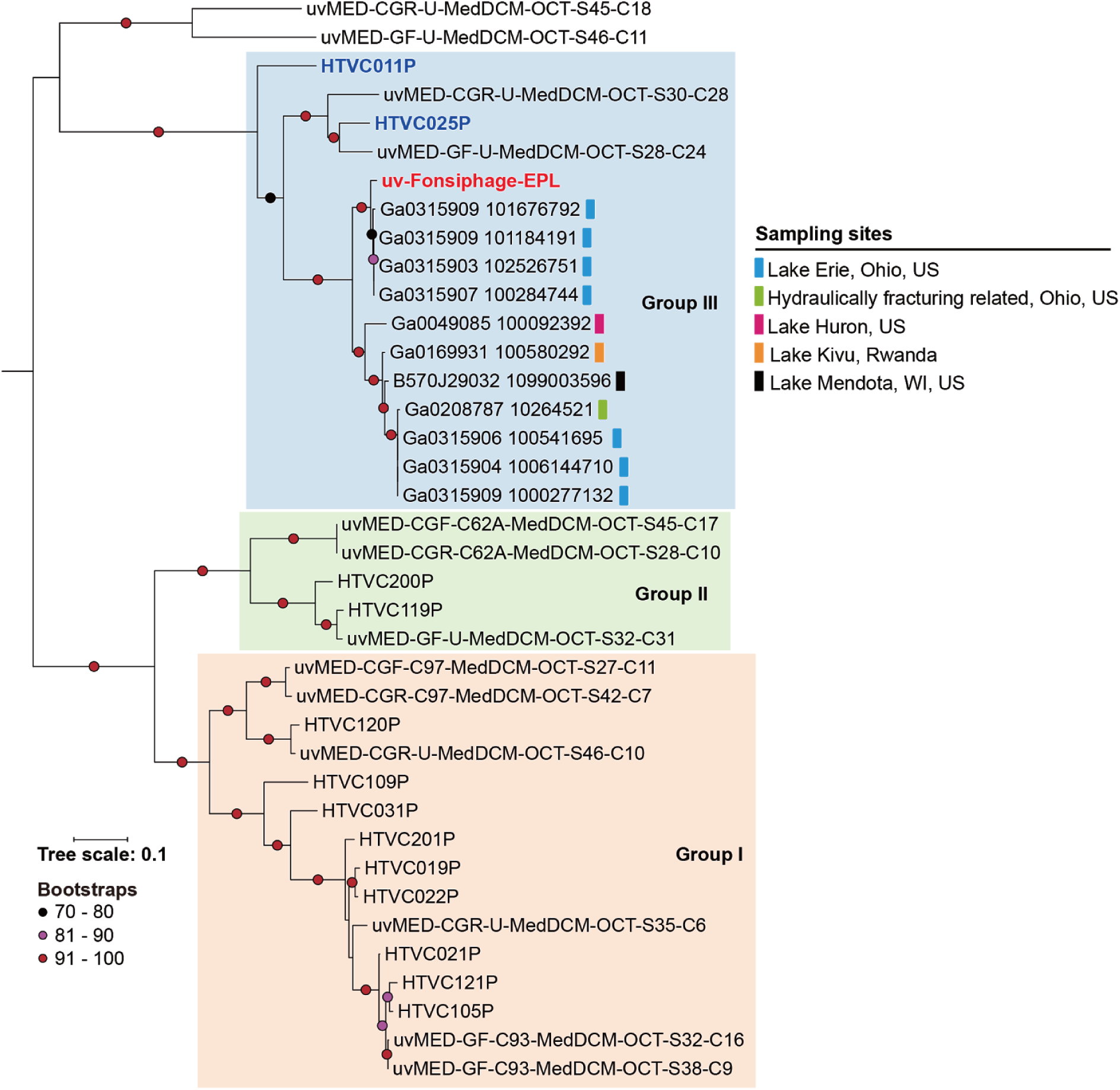
Phylogenetic analyses showed a single lineage of uv-Fonsiphage-EPL related phages detected in freshwater ecosystems. The tree was built based on the phage large terminase (TerL) proteins. Note that the TerL identified in the hydraulically fracturing related samples in Ohio was almost identical to some of those found in the Lake Erie of Ohio. The *HTVC019Pvirus* groups defined recently are shown (Zhao et al. 2018). The Pelagiphages in HTVC019Pvirus group III are indicated in blue and bold, uv-Fonsiphage-EPL is indicated in red and bold. Only one related phage TerL was from Africa (Lake Kivu), all other from North America, and it is interesting that the Africa one showed a high similarity to the one identified in Lake Mendota (98.8%), given that these two sites are quite distant from each other.

**Supplementary Fig. 11.**
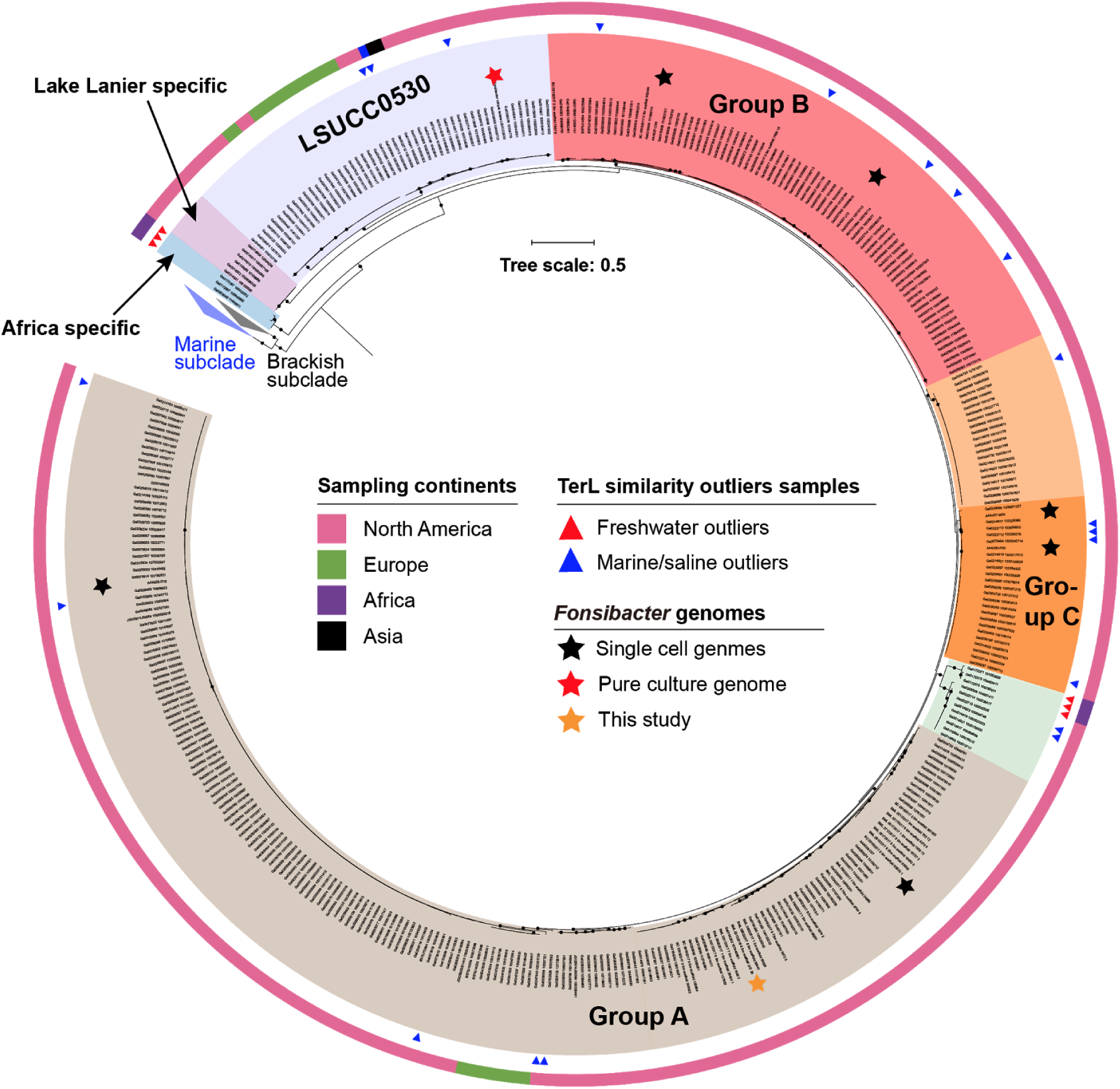
Phylogenetic analyses of *Fonsibacter* based on the ribosomal protein S3 (rpS3) nucleotide sequences. All those *Fonsibacter* rpS3 from freshwater-related samples with TerL detected, and also those from marine/saline samples with TerL similarity outliers (blue triangles), were included for analyses. *Fonsibacter* from freshwater-related samples with TerL similarity outliers are indicated by red triangles. The three groups within *Fonsibacter* are shown based on phylogeny. The nodes representing *Fonsibacter* genomes are indicated by stars. We found two lineages only detected in sampling site-specific samples, related information is shown in the left upper corner. The sampling continents are shown in color strips (the outer ring). The rpS3 sequences from marine and brackish SAR11 were used as references, and the two subclades are collapsed. The bootstraps value ≥ 90 is indicated by a black dot.

